# Quantifying subcellular organization from localization to co-occurrence with InSpace

**DOI:** 10.64898/2026.10.29.762981

**Authors:** Yiyang Niu, Ye Zheng

## Abstract

Cell identity and function depend not only on molecular abundance, but also on how molecules are organized within cells. Although spatial transcriptomics now resolves individual RNA molecules within cells, routine computational workflows reduce these measurements to cell-by-gene count matrices, retaining abundance while discarding information about intracellular localization and molecular co-occurrence. This loss of spatial information creates a need for scalable methods that preserve subcellular detail and enable comparisons across diverse cellular morphologies. Therefore, we present InSpace, a scalable framework for quantifying these complementary dimensions of subcellular organization. InSpace-R maps transcripts onto a continuous radial coordinate anchored to nuclear and cell boundaries, while InSpace-C measures distance-dependent co-occurrence or depletion of molecular pairs. Simulations and orthogonal biochemical references support accurate measurements, with scalability demonstrated on datasets containing 275 million transcripts. RNA–RNA and RNA–DNA analyses recover organelle-linked neighbourhoods and associations between nascent transcripts and their genomic origins. Analysis of B and plasma cells across 58 tumour datasets spanning 12 cancer types reveals that reduced MHC-II transcript abundance accompanies more peripheral localization and stronger mutual co-occurrence, separating these transcripts from a program enriched for ER-targeted proteins. InSpace makes intracellular organization accessible alongside abundance, enabling discovery of spatial programs that distinguish cellular states and generate hypotheses about their functional specialization.

## Background

Spatial transcriptomics now maps individual RNA molecules within cells, yet routine analyses often reduce these molecular maps to cellular gene counts, leaving a growing wealth of information about intracellular organization underused. Extracting this information requires scalable computational tools that reveal where transcripts localize, which molecules share spatial neighbourhoods, and how these patterns distinguish cellular states beyond abundance (Fig. 1a). RNA localization supports spatially restricted protein synthesis, ^1,2^ contributing to cell polarity, ^3^ migration, ^4^ developmental cell-fate decisions ^5,6^ and organelle maintenance. ^7^ These effects need not be accompanied by changes in overall abundance. For example, disruption of CPEB-dependent ZO-1 mRNA localization impairs epithelial polarity without reducing total ZO-1 protein levels. ^3^ Changes in intracellular RNA organization also accompany stress responses and contribute to tumour invasion. ^8,9^ Thus, understanding how cells regulate molecular function requires measurements of where transcripts reside and which molecules occupy shared intracellular environments.

**Figure 1:**
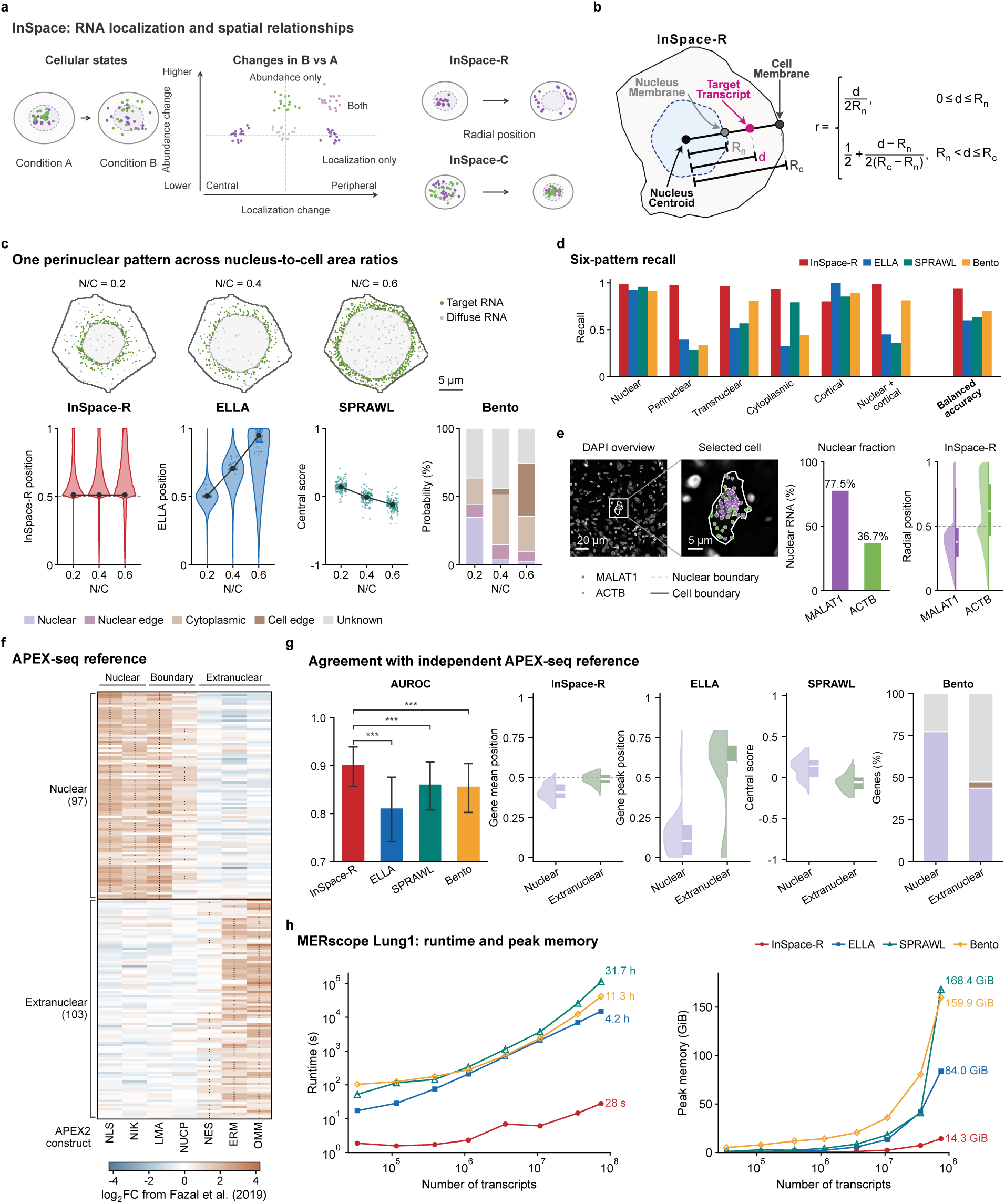
Overview and benchmarking of InSpace-R. **a**, Schematic of the InSpace framework and its two modules, InSpace-R (radial position) and InSpace-C (spatial relationships). **b**, Definition of the InSpace-R position *r*. *d* is the distance of a transcript from the nuclear centroid, and *R_n_*and *R_c_* are the distances from the centroid to the nuclear and cell boundaries along the same ray. **c**, A simulated perinuclear pattern in one cell at three nucleus-to-cell area ratios (N/C) (top), and the native readouts of InSpace-R, ELLA, SPRAWL and Bento (bottom). **d**, Recall of six simulated localization patterns and balanced accuracy for each method. **e**, MALAT1 and ACTB transcripts in a representative cell (left), and their nuclear fraction and InSpace-R positions across 12 Xenium datasets (right). **f**, log_2_ fold changes from the APEX-seq atlas of Fazal et al. for 97 nuclear and 103 extranuclear reference genes, measured with seven APEX2 constructs targeting the nucleus (NLS), nucleolus (NIK), lamina (LMA), nuclear pore (NUCP), cytosol (NES), endoplasmic reticulum membrane (ERM) and outer mitochondrial membrane (OMM). **g**, Agreement of each method with the APEX-seq reference in HEK293T Xenium data, shown as AUROC with 95% CI error bars and as per-gene readouts for nuclear and extranuclear genes. *^∗∗∗^P <* 0.001, two-sided paired bootstrap with Holm correction, InSpace-R versus each method. **h**, Runtime and peak memory of the four methods on the MERSCOPE Lung1 dataset at increasing numbers of transcripts.

Modern spatial transcriptomics maps thousands of RNA species at subcellular resolution,^10,11^ and tissue atlases now encompass millions of cells. ^12^ Multiplexed protein imaging similarly links subcellular organization to cellular states and responses to perturbation.^13,14^ Nevertheless, routine spatial workflows primarily summarize cellular abundance into cell-by-gene count matrices, leaving intracellular positioning and molecular co-occurrence underused. These measurements provide complementary information. Abundance indicates whether a molecular program is present, localization describes where its transcripts are positioned, and co-occurrence identifies shared molecular neighbourhoods. Co-occurrence can suggest common compartmental organization, although proximity alone does not establish direct molecular interaction.^15^ Systematic analysis of these spatial features could therefore reveal functional heterogeneity that abundance-focused analyses overlook.

Existing computational methods have established the feasibility and biological value of subcellular analysis. SPRAWL quantifies several localization patterns, whereas ELLA models continuous spatial variation along the cellular radius.^16,17^ Bento combines localization annotation, subcellular domain discovery and gene–gene co-localization analysis. ^18^ InSTAnT, CellSP and pyTrance further enable proximity testing, spatial module discovery and prediction of RNA co-localization. ^19–21^ A remaining challenge is to combine anatomically comparable transcript positions with interpretable co-occurrence measurements in a framework that supports large spatial datasets.

Such comparisons must account for the relationship between molecular position and cellular geometry. Broad localization labels can obscure continuous variation within compartments, while radial coordinates anchored only to the nuclear centre and cell boundary do not retain the nuclear boundary as a common anatomical reference.^17,18^ Consequently, the same normalized radius can correspond to a nuclear or cytoplasmic position in cells with different nucleus-to-cell size ratios. For co-occurrence, observed proximity must be interpreted relative to an explicit expectation that accounts for the frequencies of the molecular identities being compared. Both measurements must also remain computationally practical as molecular coverage and cell numbers increase. Addressing these challenges would support systematic investigation of how intracellular organization relates to cell identity and functional state.

Here we present InSpace, a framework for intracellular spatial position and co-occurrence enrichment. InSpace-R maps each transcript onto a continuous radial scale anchored to the nuclear centroid, nuclear boundary and cell boundary, supporting comparisons across diverse cellular geometries (Fig. 1). InSpace-C quantifies distance-dependent enrichment or depletion of molecular pairs to characterize shared intracellular neighbourhoods (Fig. 2). We evaluated both modules using simulations and orthogonal biochemical localization references, including APEX-seq, ^22^ and assessed scalability on datasets containing up to 275 million transcripts. Applications to RNA–RNA and RNA–DNA measurements demonstrate the framework’s use in describing intracellular spatial relationships. Across 58 tumour datasets spanning 12 cancer types, InSpace identifies differences in RNA localization and co-occurrence programs between B and plasma cells that complement transcript abundance. By providing interpretable spatial features for characterizing cellular phenotypes, InSpace extends spatial profiling beyond what cells express to how transcripts are organized within them and generates testable hypotheses for mechanistic studies (Fig. 3).

**Figure 2:**
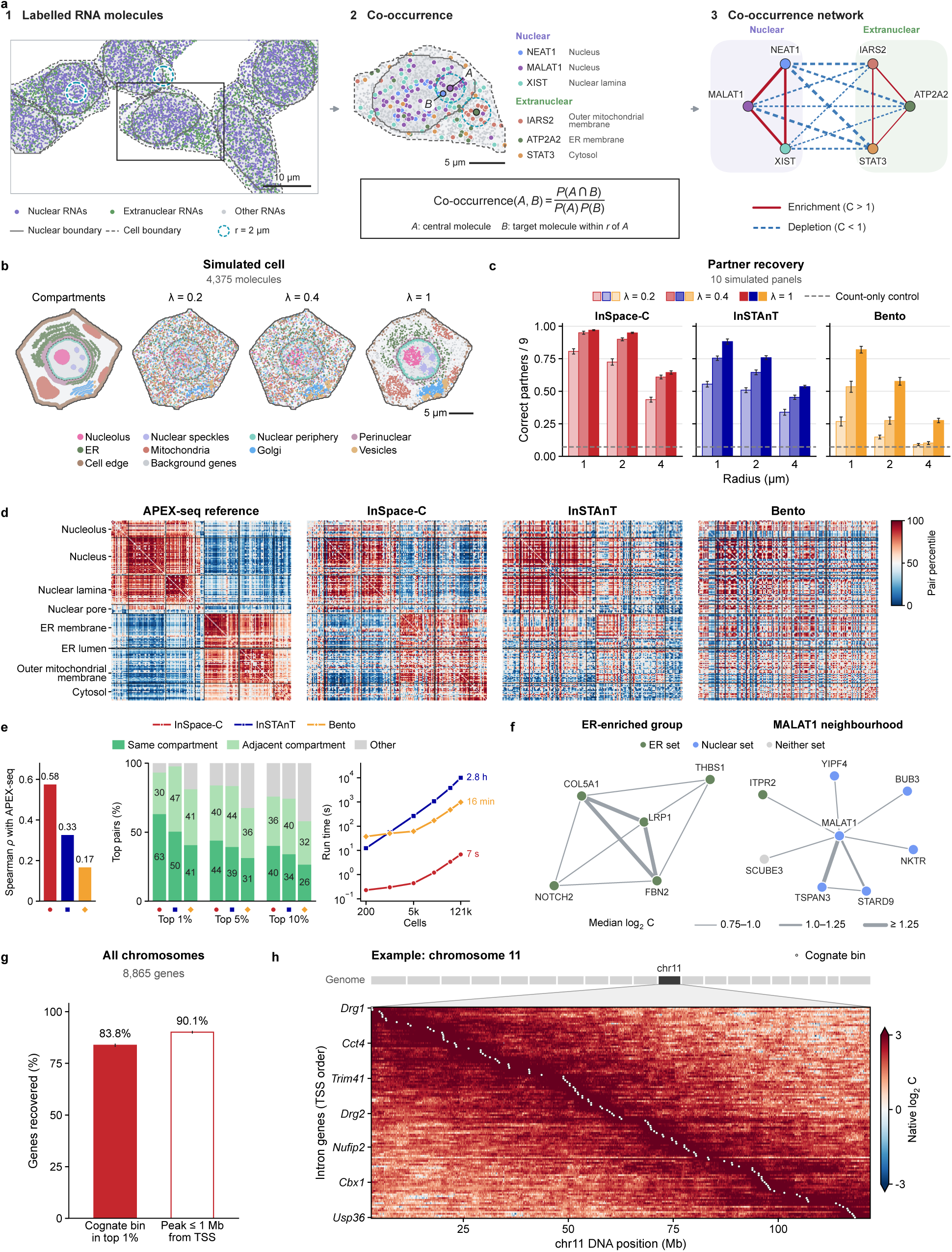
InSpace-C measures molecular co-occurrence in simulated and experimental data. **a**, Overview of InSpace-C. (1) A HEK293T Xenium field, with RNAs of the nuclear and extranuclear reference genes of Fig. 1f in purple and green, and 2-*µ*m neighbourhoods shown as circles. (2) An enlarged cell with three nuclear and three extranuclear RNAs, and the definition of co-occurrence (*C*) between a central molecule (*A*) and a target molecule within distance *r* of it (*B*). (3) Co-occurrence network of these six RNAs over the cells analysed in **d**. Red solid lines show enrichment (*C >* 1) and blue dashed lines show depletion (*C <* 1); line width scales with *|* log_2_ *C|*. **b**, A simulated cell, showing its compartments (left) and its molecules at signal fraction *λ* = 0.2, 0.4 and 1, coloured by planted module. **c**, Top-9 partner recovery in simulated data by radius and signal fraction. Bars show the mean and error bars the s.d. over *n* = 10 simulated panels; the dashed line marks the count-only control. **d**, Gene-pair co-occurrence in HEK293T Xenium data compared with APEX-seq. Matrices show percentiles of the APEX-seq profile cosine and of each method’s score over the same gene pairs, with genes ordered by APEX-seq compartment. **e**, Agreement of each method with APEX-seq on the same gene pairs. Bars show the Spearman *ρ* with the APEX-seq cosine (left) and the APEX-seq compartment relation of each method’s top 1%, 5% and 10% pairs (middle), with numbers giving the percentages of same- and adjacent-compartment pairs. Right, run time as a function of cell number. **f**, RNA co-occurrence network in U2OS MERFISH data. Edges are gene pairs above the Blank-control threshold in all four replicates, and line width gives the median log_2_ *C* over replicates in three classes. Node colour gives the RNA-localization gene set of Xia et al., ^11^ with genes in both sets assigned to the nuclear set. Node positions are schematic, and edge length carries no information. **g**, Recovery of the source neighbourhood of intron RNAs over all chromosomes: the percentage of 8,865 genes whose cognate DNA bin ranks in the top 1% (filled) or whose peak lies within 1 Mb of the transcription start site (open). Error bars show Wilson 95% confidence intervals. **h**, Example from chromosome 11: intron RNA–DNA co-occurrence for the 120 genes with the most nearby pairs, ordered by transcription start site, with cognate bins marked by white dots.

**Figure 3:**
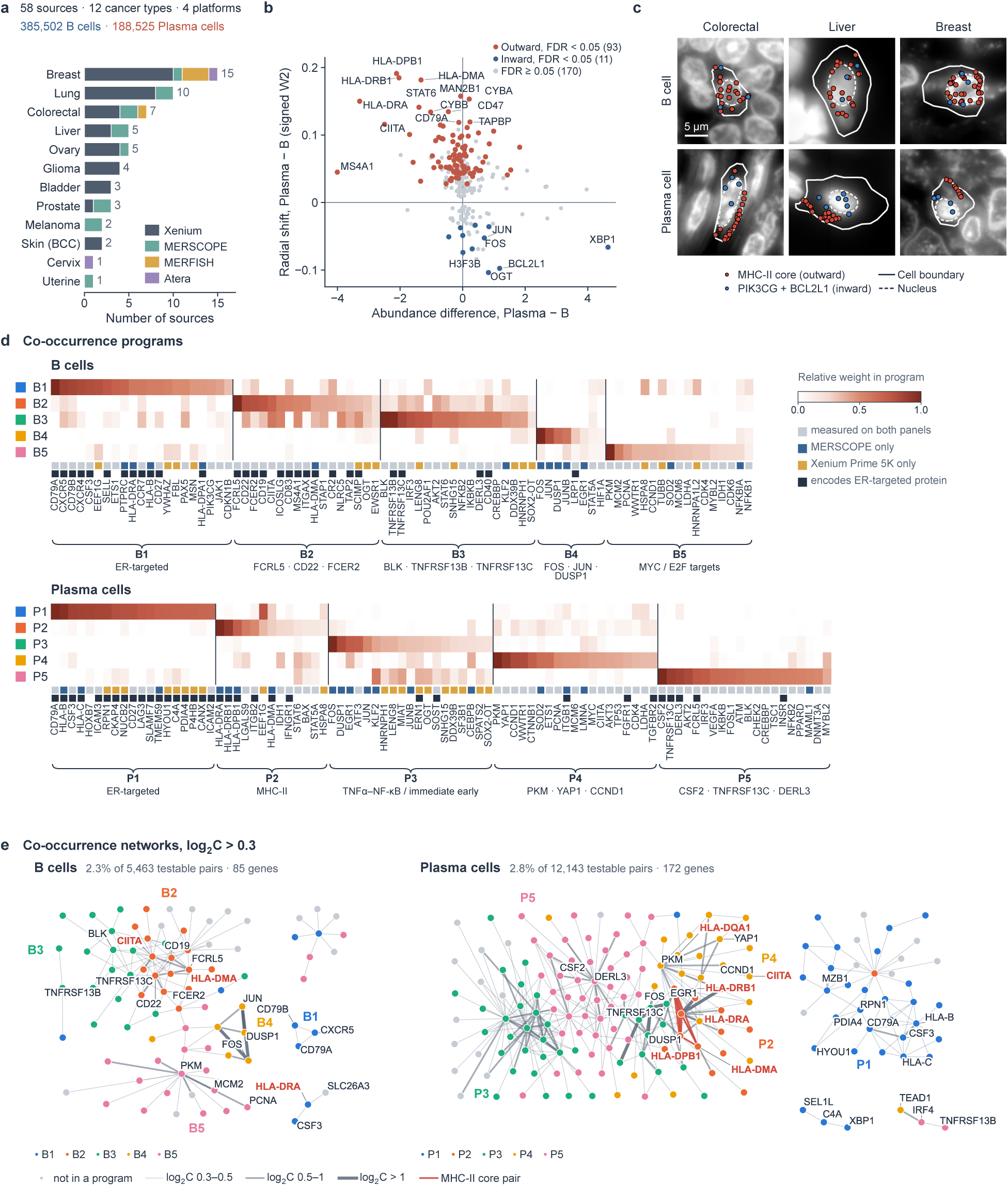
RNA radial allocation and co-localization programs in B and plasma cells. **a**, Number of sources per cancer type, coloured by platform. Header, numbers of sources, cancer types, platforms, B cells and plasma cells. **b**, Radial shift (signed W2) against abundance difference, plasma *−* B; one point per gene. Positive shift, farther from the nucleus in plasma cells. Red, outward; blue, inward (FDR *<* 0.05); grey, not significant. **c**, Example B (top) and plasma (bottom) cells on DAPI. Red, MHC-II core (*HLA-DRA*, *-DRB1*, *-DPB1*, *-DMA*); blue, *PIK3CG* and *BCL2L1*. **d**, Co-localization program weights in B cells (B1–B5) and plasma cells (P1–P5); columns, top 20 genes per program, grouped under the program holding the largest share of their weight. Colour, weight of each gene in the program relative to the program’s top gene. Squares below, measurement panel (grey, both; blue, MERSCOPE; yellow, Xenium Prime 5K) and ER-targeted protein (dark). Braces, program names (Methods). Coloured squares beside the program codes, program colours used in **e**. **e**, Co-localization networks of B cells (left) and plasma cells (right), each with its own genes. Edges, gene pairs with log_2_ *C >* 0.3; width and darkness, log_2_ *C* of 0.3–0.5, 0.5–1 or *>*1; red, pairs of MHC-II core genes. Node colour, program in **d**; grey, gene not in a program. Connected groups of fewer than three genes are not shown. Labels, the three highest-weight genes of each program, MHC-II genes (red), genes named in the text and all genes of groups of up to four genes. Header, share of testable pairs with log_2_ *C >* 0.3 and number of genes shown.

## Results

### InSpace-R recovers localization patterns across cell shapes and nuclear sizes

We compared InSpace-R (Fig. 1b and Methods) with three methods that summarize transcript position relative to cell geometry or to a fixed set of patterns: ELLA,^17^ which normalizes the distance of a transcript from the nuclear centroid by the distance to the cell boundary in the same direction and fits the resulting profile with a fixed family of 22 kernels, SPRAWL, ^16^ which scores each gene by the median rank of its transcripts among the transcripts of the same cell ordered by distance to the cell boundary or centroid, and Bento,^18^ which assigns five localization classes from 13 spatial features with a random forest trained in advance. To test whether each readout reflects the pattern rather than the morphology of the cell, we simulated seven localization patterns in 15 cells of a HEK293T Xenium dataset, ^23,24^ each at three nuclear sizes within its fixed outline (Supplementary Fig. 1, Methods and Supplementary Note 1). For a perinuclear pattern, the InSpace-R profile of the pooled transcripts peaked just outside the nuclear boundary at all three nuclear sizes (Fig. 1c). The readouts of the other methods changed although the pattern had not. The ELLA peak shifted outward from near its nuclear centre towards its membrane, the SPRAWL central score changed sign from positive to negative, and the Bento probability of the nuclear class fell from 35% to 2% while that of the cell-edge class rose to 39% (Fig. 1c and Supplementary Fig. 1c–g). In these methods, a change in nuclear geometry alone was read as a change in localization.

To quantify this over all patterns, we tested whether the full output of each method identifies the planted pattern across cell shapes. For each of the six localized patterns, all but diffuse, a template was built from the readouts of 14 cell shapes, and each gene of the held-out shape, at each nuclear size, was assigned to the nearest template (Methods). InSpace-R assigned the correct pattern with a balanced accuracy of 0.94, compared with 0.60 to 0.70 for the other methods (Fig. 1d). The difference was largest for the perinuclear pattern, which InSpace-R recognized in 98% of cases and the other methods in at most 39%, because their readouts for this pattern overlapped those of neighbouring patterns at other nuclear sizes (Supplementary Fig. 2a–d). At the largest nuclear size, InSpace-R still reached a balanced accuracy of 0.89, whereas the other methods fell below 0.6 (Supplementary Fig. 2e).

### InSpace-R separates nuclear and cytoplasmic RNAs in tissue

We next examined two transcripts of known localization in 12 Xenium sections of human breast tissue ^25,26^: MALAT1, a long non-coding RNA retained in the nucleus, and ACTB, a cytoplasmic mRNA ^27,28^ (Fig. 1e). Pooled over all sections, the distribution of MALAT1 positions lay inside the nuclear boundary and that of ACTB outside it, with medians of 0.38 and 0.62 on either side of the boundary value of 0.5. This separation matched the fractions of transcripts found within the segmented nucleus, 78% for MALAT1 and 37% for ACTB, and held in every section (Supplementary Fig. 3b). Because the nuclear boundary is a fixed landmark on the InSpace-R scale, the localization of a gene can be read directly from its distribution, without a reference distribution from the same dataset. The other methods also distinguished the two genes, each on its own scale (Supplementary Fig. 3).

### InSpace-R agrees with biochemical localization references

Segmentation-derived nuclear fractions share their boundaries with InSpace-R, so as an independent reference we used APEX-seq, which labels RNA in defined subcellular compartments of HEK293T cells by proximity biotinylation. ^22^ From the published APEX-seq enrichments alone we defined 97 nuclear and 103 extranuclear reference genes (Fig. 1f and Supplementary Fig. 4a,b) and evaluated how well each method’s gene-level readout, computed from HEK293T Xenium data ^23,24^, ranked the two groups. InSpace-R separated them with an area under the receiver operating characteristic curve (AUROC) of 0.90, compared with 0.81 to 0.86 for the other methods, and its advantage over each method was significant in paired gene bootstraps (Fig. 1g). Against a second reference, nuclear and cytoplasmic RNA-seq of fractionated mouse liver^29^ compared with liver Xenium data, ^30^ InSpace-R, SPRAWL and Bento reached AUROCs of 0.82 to 0.85 and ELLA 0.65, with InSpace-R scoring 99.6% of cell–gene pairs and Bento 57% (Supplementary Fig. 4c–h and Supplementary Note 3).

### InSpace-R scales to complete datasets

Current datasets reach hundreds of millions of transcripts and are being assembled into tissue and organism atlases. We ran the four methods on nested subsets of three such datasets, from 100 cells to all cells, under identical conditions (Methods). On the MERSCOPE human lung cancer dataset Lung1^31^, with 208,284 cells and 75 million transcripts, InSpace-R finished in 28 s with 14 GiB of peak memory, whereas ELLA, Bento and SPRAWL needed 4 to 32 h and 84 to 168 GiB (Fig. 1h). The gap widened with dataset size. InSpace-R processed the 275 million transcripts of the HEK293T Xenium data in 3 min and the 119 million transcripts of a Xenium Prime 5K lung cancer dataset^32^ in under a minute, whereas SPRAWL did not finish either dataset within the 48 h limit and Bento exhausted the 384 GiB of memory on 100,000 HEK293T cells (Supplementary Fig. 4i,j).

### InSpace-C recovers planted co-occurrence modules

InSpace-C scores how much more or less often two molecular labels lie within a chosen distance of each other than expected from their overall frequencies, pooling pairs over the cells of a population (Fig. 2a and Methods). To test whether it identifies genes that share a compartment irrespective of how their molecules are distributed within it and of how abundant they are, we simulated panels of 90 genes in nine compartment modules, placed by five different mechanisms, and 30 background genes that cluster without co-localizing with any other gene (Fig. 2b, Supplementary Fig. 5, Methods and Supplementary Note 2). For each module gene, all other genes were ranked by co-occurrence, and the fraction of its nine top-ranked partners that belonged to its module was recorded. At a radius of 2 µm, with 40% of each module gene’s molecules placed on its module and the rest spread uniformly over the cell, InSpace-C placed a true partner in 90% of these positions, compared with 65% and 27% for InSTAnT^19^ and Bento ^18^ and 7% for a control that correlated gene counts across cells (Fig. 2c). The advantage held in every simulated panel, at weaker and stronger planted signal and at radii of 1 and 4 µm (Fig. 2c), and by average precision, and when no module was planted every method remained at chance (Supplementary Fig. 6a,b).

### InSpace-C agrees with an independent localization reference

To evaluate the measure on real transcripts, we compared co-occurrence in HEK293T Xenium data ^23,24^ with the APEX-seq compartment profiles of the same cell line. ^22^ The similarity of the APEX-seq profiles of two genes served as an independent measure of their shared localization (Fig. 2d). InSpace-C scores followed this similarity with a Spearman correlation of 0.58, compared with 0.33 and 0.17 for InSTAnT and Bento, and of the 1% of pairs scored highest by InSpace-C, 63% lay in the same APEX-seq compartment, against 16% expected by chance (Fig. 2e). At the other end, 90% of the 5% of pairs that InSpace-C scored lowest lay in neither the same nor an adjacent compartment, compared with 64% and 59% for InSTAnT and Bento and 59% of all pairs (Supplementary Fig. 6c). We then ran each method on nested subsets of the same data, from 200 cells to all 121,343 cells (Methods). With 200 cells, InSpace-C already exceeded the agreement that the other methods reached on all cells (Supplementary Fig. 6d), and on the complete dataset it ran in 7 s with 19 GiB of peak memory, against 2.8 h and 178 GiB for InSTAnT and 16 min and 157 GiB for Bento (Fig. 2e and Supplementary Fig. 6e).

### InSpace-C recovers organelle-linked RNA networks

To test whether co-occurrence recovers known organization in an independent dataset without any annotation, we analysed U2OS MERFISH data with four replicates ^33^ and kept the gene pairs whose co-occurrence in all four replicates exceeded the highest value that any blank-control pair sustained across the replicates (Methods). This left 17 edges among 13 genes, which formed two groups: five genes enriched at the endoplasmic reticulum and a neighbourhood of nuclear genes around MALAT1 (Fig. 2f). Fifteen of the 17 edges joined genes that share a localization set defined in an earlier MERFISH study ^11^ (Supplementary Fig. 7). In two Cytos datasets of A549 and HeLa cells ^34,35^, the mitochondrially encoded RNAs formed a co-occurrence block of their own, and their reads lay on the mitochondrial signal (Supplementary Fig. 8 and Supplementary Note 6).

### RNA–DNA co-occurrence recovers associations with cognate genomic loci

We next extended InSpace-C from RNA alone to pairs of different molecule types. In mouse embryonic stem cells, Takei et al. imaged intron RNA and DNA loci in the same nuclei by seqFISH+^36^. We labelled each intron spot by its gene of origin and each DNA locus by its 100-kb genomic bin, and scored RNA–DNA co-occurrence within nuclei at a distance of 0.5 µm (Methods), which asks whether a nascent transcript lies near the DNA of its own gene. In single nuclei, intron RNA spots lay next to DNA loci of their own gene (Supplementary Fig. 9a). Among 8,865 genes with sufficient support, the gene’s own bin ranked in the top 1% of bins on its chromosome for 84%, and the highest-scoring bin lay within 1 Mb of the transcription start site for 90% (Fig. 2g). On each chromosome the own bin ranked in the top 1% for 69–88% of genes, lowest on chromosome 19, the shortest chromosome with the fewest candidate bins, where the peak still lay within 1 Mb for 95% (Supplementary Fig. 9b,d). Along chromosome 11, co-occurrence between intron RNAs and 100-kb DNA bins was concentrated on the diagonal, where the bin lies at the genomic position of the gene (Fig. 2h). Recovery rose with the number of pairs supporting a gene (Supplementary Fig. 9c).

### MHC-II transcripts lie farther from the nucleus in plasma cells

Plasma-cell differentiation expands the endoplasmic reticulum (ER) and extinguishes MHC class II (MHC-II) expression, ^37–40^ so we compared B and plasma cells in 58 imaging-based spatial transcriptomics sources from 12 cancer types and four platforms (385,502 B cells and 188,525 plasma cells, Fig. 3a), after removing molecules attributed to neighbouring cells (Methods and Supplementary Fig. 10a–c). As expected from this differentiation, genes of the unfolded protein response and mTORC1 signalling were more highly expressed in plasma cells, and those of allograft rejection, whose leading genes included *HLA-DRA*, *HLA-DMA* and *CD74*, in B cells (Supplementary Fig. 10d,e). Of 274 genes tested with InSpace-R, 104 differed in radial distribution between plasma and B cells (FDR *<* 0.05), and 93 of these lay farther from the nucleus in plasma cells (Fig. 3b). The three largest outward shifts were those of the MHC-II genes *HLA-DPB1*, *HLA-DRB1* and *HLA-DMA*, and *HLA-DRA* and *CIITA* also shifted outward. This was not a consequence of their lower abundance, because the MHC class I genes *HLA-B*, *HLA-C* and *TAPBP* shifted outward to a similar extent while being equally or more abundant in plasma cells. Only 11 genes shifted inward, among them *XBP1*, *BCL2L1* and *PIK3CG*. The positions of both *HLA-DRA* and *XBP1* mRNAs are known to be regulated. ^41–43^ Across sources, 92% of the per-source shifts of the detected genes, and 56 of 58 of the five outward MHC-II genes, had the cohort direction (Supplementary Fig. 11). The MHC-II shifts held in plasma cells without MHC-II-high neighbours of any cell type, and no detected gene reversed direction when the sources of any cancer type or platform were left out or used alone (Supplementary Table 2 and Supplementary Note 5).

### Co-occurrence programs separate MHC-II from ER-targeted transcripts in plasma cells

Decomposing the co-occurrence of all measured genes gave five programs in each state (Fig. 3d), supported within individual sources (191 of 193 program–source tests; Supplementary Fig. 12a,b); one B-cell program, of immediate-early transcripts (B4), had no stable members. In B cells, one program (B1) joined B-cell receptor transcripts with MHC-I and MHC-II transcripts, mirroring the proximity of MHC-II to the B-cell receptor protein,^44^ and 58% of its members encode ER-targeted proteins (35% of all genes; *q* = 1.5 *×* 10*^−^*^4^). In plasma cells, MHC-II transcripts formed their own program (P2; MHC class II protein complex assembly, *q* = 0.011), whereas MHC-I transcripts joined *MZB1* and ER protein-processing transcripts such as *RPN1*, *PDIA4* and *HYOU1* (P1), 60% of whose members encode ER-targeted proteins, against 37% of all genes (*q* = 2 *×* 10*^−^*^8^). Because MHC-II chains are also translated into the ER,^45^ this separation is not one between ER-associated and cytosolic RNAs. Independent measurements place several P1 transcripts at the ER^11,46^ (Supplementary Table 1). MHC-II transcripts also lay closer to one another in plasma than in B cells: all 11 testable pairs of MHC-II module genes co-occurred more within 2 µm (mean increase in log_2_ *C* of 1.04, permutation *P ≤* 2 *×* 10*^−^*^5^). Among the pairs with log_2_ *C >* 0.3, the two strongest in plasma cells were *HLA-DRA*–*HLA-DRB1* and *HLA-DRA*–*HLA-DPB1* (log_2_ *C* of 2.36 and 1.84), and the MHC-II network was separate from that of P1, whereas no MHC-II pair reached this level in B cells, in which *CIITA* instead linked to B-cell identity transcripts such as *CD19*, *CD22* and *FCRL5* (Fig. 3e). The inward genes *PIK3CG* and *BCL2L1* co-occurred less with the MHC-II core, so that in plasma cells the MHC-II core lies peripherally and in close mutual proximity while these transcripts lie towards the nucleus (Fig. 3c). MHC-II mRNAs share a ribonucleoprotein complex that coordinates their levels, ^47,48^ a candidate basis for this organization. With the programs fixed, the genes of six held-out TNBC lymph-node sources, profiled on a panel without HLA genes, were placed on the same programs for the eight programs with enough genes on that panel (correlation of gene weights 0.69–0.92 for B1–B3, B5 and P1, 0.47–0.56 for P2 and P5, and 0.14 for P4; all *q <* 0.05; Supplementary Fig. 12c,d).

## Discussion

InSpace adds intracellular organization to the molecular features that can be compared across cell populations. InSpace-R provides a continuous description of transcript position relative to nuclear and cell boundaries, whereas InSpace-C measures enrichment or depletion of molecular identities among nearby pairs. Together, these readouts support comparisons of transcript distributions, molecular neighbourhoods and spatial programs alongside differential abundance. Their computational efficiency makes these analyses practical across complete datasets, allowing existing spatial measurements to be revisited for biological information that is lost when transcript coordinates are reduced to cellular counts. The RNA–DNA application further demonstrates that the co-occurrence framework can accommodate different types of jointly imaged molecular labels.

The B- and plasma-cell analysis illustrates why localization and co-occurrence should be considered together. Reduced expression does not necessarily imply a loss of spatial organization among the remaining transcripts, and similar radial shifts need not correspond to membership in the same molecular neighbourhood. These observations suggest that cellular specialization can involve selective reorganization of transcripts in addition to changes in their abundance. The separation of the MHC-II program from a program enriched for transcripts encoding ER-targeted proteins motivates investigation of RNA sorting, retention and translation within the secretory system. These data do not resolve the precise organelle association of these transcripts or establish that their spatial organization directly changes antigen presentation. Joint imaging and targeted perturbations could test these possibilities.

Several features of the measurements delimit their interpretation. Boundary anchoring improves anatomical comparability but does not remove all effects of cell morphology or available compartment area. Radial profiles also compress directional information, and two-dimensional proximity cannot fully describe three-dimensional organization. InSpace-C measures population-level enrichment among observed neighbouring pairs. Its reference depends on the measured molecular panel and the cells contributing those pairs, and shared compartment occupancy or cellular heterogeneity can contribute to enrichment. Co-occurrence therefore describes a spatial relationship rather than direct molecular interaction.^15^ Segmentation errors, overlapping cells and incomplete molecular detection remain additional sources of uncertainty. Future developments should incorporate geometry-aware reference distributions, compartment-conditioned co-occurrence models and hierarchical analyses that retain cell and donor variability. These extensions would help distinguish broad changes in intracellular architecture from more specific molecular associations.

An important next direction is continuous subcellular localization analysis for spatial proteomics. Multiplexed imaging and biochemical organelle profiling have shown that protein redistribution can reveal cellular states and perturbation responses that are incompletely captured by total abundance. ^13,14,49^ Extending the anatomical coordinate system to protein images would require intensity-aware models that account for imaging background, regional area and overall signal strength. Such profiles could quantify redistribution within and between nuclear, cytoplasmic and membrane-proximal regions, providing complementary phenotypes for studying signalling, organelle remodelling and treatment responses. This would build on existing protein-imaging approaches while enabling direct comparison with RNA localization on a common spatial reference.

Protein–protein and RNA–protein co-occurrence offer a further opportunity to connect molecular organization across modalities. Distance-dependent, normalized joint protein intensity could describe protein neighbourhoods, while protein enrichment around individual RNA coordinates could characterize the environments occupied by transcripts. These measurements could test whether cell-state changes involve altered association with RNA-processing compartments, translation machinery or organelle markers. Joint RNA and protein imaging has already revealed complex relationships between transcript targeting and protein production, ^50^ making this a biologically motivated extension. However, RNA–protein proximity alone would not establish local translation or binding. Such conclusions would require measurements of nascent protein synthesis, molecular interaction or responses to targeted perturbation.

These extensions require validation suited to each modality, including accurate channel registration, spatial scales matched to effective optical resolution and reference models that distinguish shared compartment preference from additional local association. Protein measurements must also account for antibody specificity, background and signal spillover. With these controls, integrating abundance, localization and molecular neighbourhoods could support a more complete description of cell identity, functional heterogeneity and adaptation. Combined with perturbation experiments and tissue context, these features could help identify which aspects of intracellular organization are conserved across cell types and which change with environmental or therapeutic pressures.

## Conclusions

InSpace provides a practical foundation for comparing intracellular organization alongside molecular abundance. Its extension across RNA and protein measurements could connect spatial phenotypes to the processes that establish and maintain cellular function. The broader opportunity is to understand how cells organize a shared molecular repertoire into distinct functional states.

## Methods

### InSpace-R radial coordinate

InSpace-R places each transcript on a radial scale anchored to its own cell and nucleus (Fig. 1b). Along the ray from the centroid of the nuclear polygon through the transcript, *d* is the distance of the transcript from the centroid, and *R_n_* and *R_c_* are the distances to the nuclear and cell boundaries. The radial position is

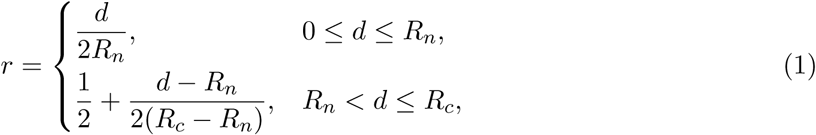

so that *r* = 0, 0.5 and 1 mark the nuclear centroid, the nuclear boundary and the cell boundary in every cell, whatever its size or shape. In concave cells, where a ray can cross the cell boundary more than once, *R_c_* is the first crossing beyond the transcript. The InSpace-R position of a gene is the mean *r* of its transcripts.

### Data quality control

Every dataset passed the same quality control before any method was run. Cells were taken from the quality-controlled cell list of each dataset and required a valid cell polygon and a valid nucleus polygon, with any nucleus extending beyond the cell outline clipped to it. Transcripts were required to fall inside their assigned cell polygon, negative-control and blank features were removed, and vendor quality thresholds were applied where available (Xenium QV *≥* 20, MERSCOPE QV *≥* 0.95). Method outputs that carried no prediction were excluded from gene-level summaries.

### Comparator methods

We compared InSpace-R with ELLA v1.0.0, ^17^ SPRAWL v0.5^16^ and Bento v2.1.4.post2. ^18^ Each method has a single-value readout, used in the gene-level comparisons, and a full output, used in the simulated benchmark. For InSpace-R these are the gene mean position and the 99 quantiles of the gene’s positions. ELLA fits a non-homogeneous Poisson process along its own radial coordinate (100 angular sectors, 22 kernels), its readout is the peak of the fitted intensity, and its full output is the fitted intensity curve (21 standardized values and their 20 differences). SPRAWL scores each cell–gene pair against permuted transcript positions on four metrics (peripheral, central, radial and punctate), its readout is the central score, which is positive when transcripts lie closer to the cell centre than expected, and its full output is the median of each metric over cells. Bento’s RNAforest classifier gives each cell–gene pair probabilities for five classes (nuclear, nuclear edge, cytoplasmic, cell edge and none), computed here from all trees of the classifier after correcting the wrapper, which scored only the first tree. Its readout is a contrast of these probabilities, defined below, and its full output is their mean over cells.

### Simulated localization benchmark

Simulated cells were built from 15 cells of the HEK293T Xenium data (Supplementary Fig. 1a), chosen to span the shapes of that dataset (Supplementary Note 1). Each cell was set to nucleus-to-cell area ratios (N/C) of 0.2, 0.4 and 0.6 by scaling its own nucleus about the nuclear centroid within the fixed cell outline. Transcripts of 70 genes followed seven patterns with ten genes each (Supplementary Fig. 1b and Supplementary Note 1): diffuse, nuclear, perinuclear, transnuclear, cytoplasmic, cortical and nuclear + cortical. Ten cells were sampled for each shape and N/C level.

The unit of analysis is a gene in one shape at one N/C level, pooling its transcripts over the ten cells (2,700 units with a localized pattern), described by the full output of its method (Comparator methods). Each unit was assigned to the nearest pattern template, the mean feature vector of each pattern over the other 14 shapes (leave-one-shape-out, root-mean-square distance). Balanced accuracy (BA) is the mean recall over the six patterns, with 95% intervals from 2,000 bootstrap resamples of the 15 shapes.

### MALAT1 and ACTB in breast tissue Xenium data

We analysed the 12 sections of the 10x Genomics Xenium FFPE human breast dataset^25,26^ with the vendor segmentation, using 66,843 cells sampled across N/C levels. The nuclear fraction of a gene is the fraction of its transcripts inside the nuclear polygon. InSpace-R positions were pooled over all cells, and ELLA, SPRAWL and Bento were run on the same cells (Supplementary Fig. 3).

### Agreement with the APEX-seq reference

HEK293T Xenium data were from the SubCellSpace study, ^23,24^ whose 290-gene panel was designed from the APEX-seq atlas.^22^ Of these genes, 250 had APEX-seq annotation (Supplementary Fig. 4a,b). A gene was enriched in a compartment when its published log_2_ fold change exceeded 0.75 at an adjusted *P* below 0.05, the thresholds of the atlas. Nuclear reference genes were enriched in the nucleus (NLS) or nucleolus (NIK) and in no extranuclear compartment. Extranuclear reference genes were enriched in the cytosol (NES), endoplasmic reticulum membrane (ERM) or outer mitochondrial membrane (OMM) and in no nuclear or nuclear-boundary compartment (lamina, LMA, and nuclear pore, NUCP). Enrichment in at least one of these compartments also had to hold in both replicates. This gave 97 nuclear and 103 extranuclear reference genes. We used 8,000 randomly sampled quality-controlled HEK293T cells. Gene readouts were as under Comparator methods, with the Bento contrast the nuclear and nuclear-edge probabilities minus the cytoplasmic and cell-edge probabilities. Agreement with the reference genes was measured by the area under the receiver operating characteristic curve (AUROC), with InSpace-R and ELLA positions negated so that higher values indicate nuclear localization. 95% CIs and paired differences from InSpace-R were obtained from 20,000 gene bootstrap resamples drawn within each reference group. Two-sided *P* values for these differences were taken from the same resamples and adjusted for the three comparisons with the Holm method (Fig. 1g).

### Runtime and memory benchmark

Runtime and peak memory were measured on the MERSCOPE human lung cancer dataset Lung1^31^ (208,284 cells, 75 million transcripts), the HEK293T Xenium data (121,343 cells, 275 million transcripts) and the Xenium Prime 5K human lung cancer dataset^32^ (260,617 cells, 119 million transcripts), at eight nested cell subsets from 100 cells to the full dataset. Each method ran once per subset in a fresh container with 32 CPU cores and 384 GiB of memory, with ELLA at its default 360 angular sectors. Runtime is the wall-clock time from launch to written outputs, and peak memory is the peak memory use of the container. Runs were limited to 48 h (Fig. 1h and Supplementary Fig. 4i,j).

### InSpace-C co-occurrence for labelled molecules

InSpace-C treats each detected molecule as a point with planar coordinates and a discrete label, such as the gene of a transcript, the gene of origin of an intron spot or the genomic bin of a DNA locus (Fig. 2a). It considers all ordered pairs of distinct molecules in the same cell that lie within a chosen distance of each other. For labels *a* and *b*, let *A* be the event that a pair drawn uniformly from these pairs carries label *a* at its first molecule, and *B* the event that it carries label *b* at its second. The co-occurrence score is

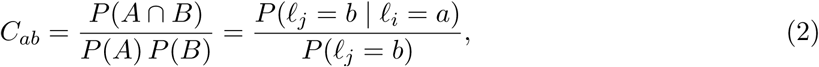

where *ℓ_i_* and *ℓ_j_* are the labels of the first and second molecule of the pair. Values above 1 indicate enrichment and values below 1 depletion, and we report log_2_ *C_ab_*. Pairs are counted over all cells of a population before the probabilities are formed, every counted label enters the denominator, and the score is symmetric in *a* and *b*.

### Simulated co-occurrence benchmark

Each simulated panel comprised 150 cells, ten from each of 15 HEK293T cell shapes at a nucleus-to-cell area ratio of 0.2 with nine compartment regions, and 120 genes: 90 module genes, two for each combination of nine compartment modules and five placement mechanisms, and 30 background genes that cluster without co-localizing with any other gene (Supplementary Fig. 5 and Supplementary Note 2). Abundances were assigned independently of module membership. Each molecule of a module gene was placed on its module with probability *λ* and uniformly over the cell otherwise, and ten panels were generated at each *λ ∈ {*0, 0.2, 0.4, 0.6, 0.8, 1*}*. InSpace-C, InSTAnT^19^ (sc-instant 1.2.1), Bento^18^ and a count-only control, the Spearman correlation of gene counts across cells, were run per cell at a radius of 2 µm, with 1 and 4 µm as sensitivity analyses (Supplementary Note 2). Scores were rank-normalized over all gene pairs, with unscored pairs at the median rank. Each module gene served as a query, with its nine module partners as positives and the other 119 genes as candidates. The top-9 hit rate is the fraction of a query’s nine highest-ranked candidates that belong to its module, averaged over queries (9/119 at random), and average precision was averaged over queries. Both are reported as mean *±* s.d. over the ten panels at each *λ* and radius, with methods compared by paired per-panel differences and 95% confidence intervals (Fig. 2c and Supplementary Fig. 6a,b).

### Agreement with APEX-seq and scaling in HEK293T Xenium data

The HEK293T Xenium data were the 121,343 quality-controlled cells and 290 genes of the Figure 1 runtime benchmark, scored by InSpace-C, InSTAnT and Bento at 2 µm (Supplementary Note 4). The reference was the APEX-seq^22^ enrichment profile of each of the 250 annotated panel genes across eight compartments; the reference for a gene pair was the cosine of the two centred profiles, and pairs were classified as same compartment, adjacent compartment or other from the compartment of maximal enrichment of each gene (Supplementary Note 4). Comparisons used the 13,299 pairs of 178 genes scored by all three methods: the Spearman correlation of each method’s scores with the APEX-seq cosine, and the compartment relation of its top and bottom 1%, 5% and 10% pairs (Fig. 2d,e and Supplementary Fig. 6c). A secondary analysis ranked each method within all pairs it scored and compared its top pairs with 5,000 gene-label permutations of the reference. For scaling, each method ran on nested random subsets of 200 to 60,000 cells and on all cells, one job per run on a node with 48 CPU cores and 384 GiB of memory (Fig. 2e and Supplementary Fig. 6d). Run time was the wall-clock time of the method call, excluding input loading and output writing, and peak memory the peak resident set size of the method process (Supplementary Fig. 6e).

### U2OS MERFISH co-occurrence network

InSpace-C was run at 2 µm on each of four replicates of the public U2OS MERFISH dataset ^33^ (16,729 cells, 130 genes and 10 Blank controls). A gene pair was kept as an edge when, in every replicate, it had at least 20 observed pairs, at least 20 pairs expected under independence and log_2_ *C* above 0.665. This threshold is the highest log_2_ *C* that any pair involving a Blank control reached in all four replicates under the same count conditions. Edge width in Fig. 2f shows the median log_2_ *C* over replicates in three classes. Nodes are coloured by membership of the ER-enriched (log_2_ fold change *>* 0) and nuclear-enriched (log_2_ fold change *>* 2) gene sets of Xia et al., ^11^ both at Bonferroni-adjusted *P <* 10*^−10^*; a gene in both sets was assigned to the nuclear set, because the ER test of Xia et al. compares only cytoplasmic molecules (Supplementary Fig. 7).

### RNA–DNA co-occurrence

Intron RNA and DNA seqFISH data from mouse embryonic stem cells (E14; two biological replicates) ^36^ were scored in two dimensions within nuclei over [0, 0.5) µm, with intron spots labelled by gene of origin, DNA loci by 100-kb genomic bin, all RNA–RNA, RNA–DNA and DNA–DNA pairs in the background, and counts pooled over replicates. For each gene, candidates were the scorable bins on its own chromosome; the cognate bin was counted as recovered when it ranked in the top 1% of candidates, and the peak, the highest-scoring candidate bin, when it lay within 1 Mb of the transcription start site (Fig. 2g,h). Recovery was evaluated on the 8,865 genes with at least 2,000 same-chromosome intron–DNA pairs and a scorable cognate bin, overall and per chromosome (Supplementary Fig. 9 and Supplementary Note 7).

### B and plasma cell cohort

The cohort comprised 58 human tumour sources from 12 cancer types, profiled on Xenium and Atera (10x Genomics), MERSCOPE and MERFISH, the Xenium sources including Xenium Prime 5K panels, with 385,502 B cells and 188,525 plasma cells (Fig. 3a and Supplementary Note 5). Each source was the unit of inference. B and plasma cells were taken from harmonized B-lineage labels, obtained by reclustering each dataset at three hierarchical tiers and reviewing every cluster against canonical markers and differentially expressed genes, excluding ambiguous cells, and passed the filters under Data quality control. Molecules attributed to other cell types were removed with cellAdmix^51^ (v0.1.0) in marker mode: within each source, molecular neighbourhoods were factorized, each molecule of a B or plasma cell was assigned to a factor, and molecules of factors enriched for markers of other lineages were removed (Supplementary Fig. 10a–c and Supplementary Note 5). The genes under study (HLA class II genes, *CD74* and *CIITA*) and *XBP1* and *IRF4* were never markers. HLA class II genes other than *HLA-DRA* and *CIITA* were on MERSCOPE panels only, and the Xenium Prime 5K panels carried no HLA genes.

### Differential radial localization

Radial distributions were compared between plasma and B cells with the across-source test of InSpace-R. For each source *s*, 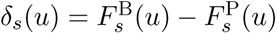 is the contrast between the empirical distribution functions of *r* over the transcripts of each state, evaluated at 199 midpoints of [0, 1], and with *k* sources weighted equally the statistic is the integrated squared mean contrast,

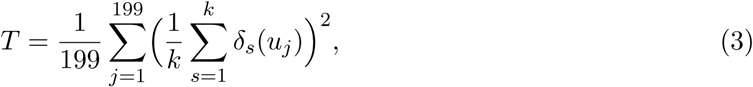

Its null distribution reverses the signs of whole source contrasts. Genes entering at least six sources were tested, with Benjamini–Hochberg adjustment^52^ (Fig. 3b and Supplementary Note 5). The effect size is the 2-Wasserstein distance *W*_2_ between the equal-weight mixtures of the source distributions of each state, signed so that positive values indicate transcripts farther from the nucleus in plasma cells, and the abundance difference is the mean log-normalized count in plasma minus B cells, averaged over sources. Gene sets that differed in expression were identified by preranked GSEA of the Hallmark sets, with genes ranked by a one-sample test of their source differences (Supplementary Fig. 10d,e and Supplementary Note 5).

### Differential co-occurrence

For each source and state, InSpace-C co-occurrence *C_ab_* within 2 µm was computed for all gene pairs of the panel, without gene selection, over all cells, with all retained molecules as the background, and a pair was supported in a source and state when it met fixed cell-number and count thresholds (Supplementary Note 5). Differences were tested in the 16 MERSCOPE or MERFISH sources. For each pair, *D* is the mean over sources supporting the pair in both states of log_2_ *C*^P^ *−* log_2_ *C*^B^, and pairs supported in at least three sources were tested against 100,000 permutations of the B and plasma labels among cells within each source, with two-sided *P* values adjusted by the Benjamini– Hochberg procedure (Fig. 3c). The MHC-II module statistic, the mean *D* over pairs among 11 predefined MHC-II genes, was tested with the same permutations. These *P* values refer to cells within the sampled sources. As neighbour controls, both tests were repeated on plasma cells with no MHC-II-high cell of any type within 10, 15 or 20 µm and on B cells with no such cell outside the B lineage, a neighbour being MHC-II-high when its HLA class II transcript count reached the median of the plasma cells in its source (Supplementary Table 2 and Supplementary Note 5). The co-occurrence test, the neighbour controls and the TNBC replication followed plans fixed before analysis.

### Co-occurrence programs

Programs were identified separately in B and plasma cells from 16 MERSCOPE or MERFISH and 18 Xenium Prime 5K sources, with the six TNBC lymph-node sources held out. For each state, the matrix *A_ab_* = max(median*_s_* log_2_ *C_ab_,* 0) over pairs supported in at least six sources, with other entries masked, was approximated as *HH^⊤^* with non-negative *H* by symmetric NMF^53,54^ fitted to observed entries (171 genes in B cells, 340 in plasma cells). The number of programs was chosen by held-out error (five in each state), consensus gene weights (NMF loadings) were medians over 50 random starts, and stable members were genes whose weight fell consistently on one program across starts (Supplementary Note 5). Fig. 3d shows the union of the 20 highest-weight genes of each program, relative to the program’s top gene, with ER-targeted proteins annotated from UniProtKB/Swiss-Prot^55^. Stable members of each program were tested against all genes of the state’s fit for enrichment of ER-targeted proteins and of MSigDB Hallmark (v2026.1)^56^ and Gene Ontology Biological Process (2025)^57^ sets (one-sided hypergeometric test, Benjamini–Hochberg adjustment per program and collection). Programs with an enriched set at *q <* 0.05 were named after it (B1 and P1, ER-targeted; B5, MYC and E2F targets; P2, MHC-II, after MHC class II protein complex assembly; P3, TNF*α*–NF-*κ*B, whose leading genes are immediate-early genes) and the others after their three highest-weight genes; the enriched sets of P4 were general transcription terms, and B4, without stable members, was named after its three highest-weight genes. Fig. 3e shows, for each state separately, the pairs supported in at least six sources with median log_2_ *C >* 0.3 (the same threshold in both states), in connected groups of at least three genes, with each gene coloured by the program holding the largest share of its weight. For replication in the held-out TNBC sources, the discovery programs were kept fixed. A program was recoverable when at least five of its stable members and ten pairs among them were testable on the TNBC panel, and each TNBC gene was placed on the fixed programs by non-negative least squares over its testable partners. A program replicated when the correlation between these TNBC weights and the discovery weights exceeded that under 10,000 permutations of the TNBC gene labels (*q <* 0.05, Benjamini–Hochberg; Supplementary Fig. 12c,d and Supplementary Note 5).

### Software and statistics

InSpace is implemented in the Python package inspace (v1.0.0) with a Rust core. Analyses used Python 3.11 with NumPy 2.5.2^58^, pandas 3.0.5^59^, SciPy 1.18.1^60^ and scikit-learn 1.9.0^61^, and cellAdmix 0.1.0 in R 4.5.1. Figures were drawn with Matplotlib 3.10.9^62^ and CNSPlots 0.6.0^63^. Bootstrap intervals are percentile intervals.

## Data availability

This study reanalysed publicly available datasets. No new experimental data were generated.

Processed HEK293T Xenium validation data from the SubCellSpace study ^23,24^ are available from Zenodo at https://zenodo.org/records/19708995. The APEX-seq subcellular RNA-localization atlas ^22^ is available from the Gene Expression Omnibus (GEO) under accession GSE116008.

The 12 human breast FFPE Xenium sections used for MALAT1 and ACTB localization benchmarking ^25,26^ are available from https://www.10xgenomics.com/datasets/xenium-ffpe-human-breast-biomarkers. The Xenium Prime 5K human lung cancer dataset used for runtime and memory benchmarking ^32^ is available as Experiment 2 at https://www.10xgenomics.com/datasets/xenium-human-lung-cancer-post-xenium-technote. The MERSCOPE Lung1 dataset (HumanLungCancerPatient1)^31^ is available through the Vizgen MERFISH FFPE Human Immuno-oncology Data Set at https://info.vizgen.com/ffpe-showcase. Public Xenium and Atera datasets can be accessed through the 10x Genomics dataset portal at https://www.10xgenomics.com/datasets.

Mouse liver Xenium Prime data ^30^ are available under GEO accession GSE310392; the samples analysed were GSM9295277, GSM9295280, GSM9295283 and GSM9295287. Nuclear and cytoplasmic fractionation RNA-seq data from mouse liver^29^ are available under GSE73977.

The U2OS MERFISH data of Moffitt et al. ^33^, from which four replicates were analysed, are available from http://zhuang.harvard.edu/MERFISHData/data_for_release.zip. Nuclear and endoplasmic-reticulum RNA-localization reference measurements from Xia et al. ^11^ are available in the supplementary datasets accompanying https://doi.org/10.1073/pnas.1912459116, including Dataset S4 for endoplasmic-reticulum enrichment.

The E14 mouse embryonic stem-cell DNA seqFISH+ and intron RNA imaging data ^36^ are available from https://doi.org/10.5281/zenodo.7693825. The deposit includes E14 rep1.tar.gz, E14 rep2.tar.gz and transcriptomic-data.zip, containing DNA-locus and RNA-spot coordinates. Additional processed data and annotation resources are available at https://github.com/CaiGroup/dna-seqfish-plus-multi-omics.

The Cytos A549 pooled-screening data ^34^ are available as the MBA-0832 release; vehicle-treated wells carrying non-targeting guides were analysed. The Cytos HeLa tunicamycin-response data ^35^ are available from https://www.elementbiosciences.com/resources/dataset-profiling-compound-response-of-tunicamycin-with-cytos; DMSO-treated wells from Cytos-3Prime HeLa-Tunicamycin, Run2 WTA 3, were analysed.

RNA-seq measurements of transcripts associated with LRRC59- and SEC61*β*-labelled ribosomes ^46^ are available in the supporting information accompanying https://doi.org/10.1074/jbc.RA119.007996. Protein annotations were obtained from UniProtKB/Swiss-Prot^55^, release 2026 03 (https://www.uniprot.org/). Gene-set analyses used the MSigDB Hallmark collection ^56^, version 2026.1 (https://www.gsea-msigdb.org/gsea/msigdb/human/collections.jsp), and Gene Ontology Biological Process gene sets^57^ (2025; https://geneontology.org/). Lineage-marker annotations were compiled from the published atlases and databases cited in Supplementary Note 5.

## Code availability

Custom code used for data processing, modelling, statistical analysis, and figure generation will be made available at https://github.com/CompBioWizard/InSpace. The repository includes the analysis scripts and documentation required to reproduce the computational analyses and figures presented in this study.

## Acknowledgments

This work was supported by the National Institutes of Health NIH/NHGRI grant (R00HG012797), CCSG New Faculty Award supported by the NIH/NCI grant (P30CA016672), and start-up funds from the University of Texas MD Anderson Cancer Center to Y.Z.

## Author Contributions

Y.N. and Y.Z. conceptualized the project. Under the supervision of Y.Z., Y.N. designed the study, performed all analyses and validation, and drafted the manuscript. Y.Z. reviewed and edited the manuscript.

