## Supplementary Notes and Figures for "Quantifying subcellular organization from localization to co-occurrence with InSpace"

**Supplementary Note 1. Planted localization patterns**

The 15 cell shapes of the simulated localization benchmark are cells of the HEK293T Xenium data.[1,2] Candidates were 30,000 randomly sampled analysis cells. A cell was eligible when its area lay between the 10th and 90th percentiles of the sample, its outline had no horizontal or vertical straight stretch longer than 2.5 $\mu$m and no protrusion thinner than 2 $\mu$m, which mark crop edges and segmentation spurs, and its own nucleus, scaled about its centroid to an N/C of 0.6, stayed inside the cell with at least 0.4 $\mu$m of cytoplasm, about two pixels of the segmentation masks, so that a cytoplasm remains at every nuclear size. Of the 99 eligible cells, the 14 nearest the centres of 14 $k$-means clusters of aspect ratio, solidity, circularity and area were taken (H02–H15). The example cell H01 in Fig. 1c and Supplementary Fig. 1b–g is the round eligible cell (aspect ratio below 1.25, circularity at least 0.85) with the widest cytoplasm at an N/C of 0.6.

In the simulated localization benchmark (Methods and Supplementary Fig. 1), perinuclear and transnuclear densities decayed exponentially from the nuclear boundary (decay length 0.70 $\mu$m, outside it or on both sides), the cortical density decayed from the cell boundary (0.80 $\mu$m), and the other patterns were uniform over their compartments. Transcript counts per cell were Poisson with gene means of 2.8–123 transcripts, and a gene-specific fraction of 0.65–0.90 of each gene's transcripts followed its pattern while the rest were diffuse.

**Supplementary Note 2. Simulated co-occurrence benchmark**

**Cells and genes.** The 15 cell shapes were the HEK293T cells H01–H15 of Supplementary Note 1 with the nucleus at a nucleus-to-cell area ratio of 0.2 (cell areas 178–293 $\mu m^2$). Regions for the nucleolus, nuclear speckles, endoplasmic reticulum (ER), mitochondria, Golgi and vesicles were transferred from 15 hand-drawn reference cells by a nucleus-centred map that preserves angle and relative depth between the nuclear and cell boundaries, and were restricted to the nucleus or cytoplasm of the target cell. The nuclear periphery (within 0.4 $\mu$m inside the nucleus), the perinuclear region (within 1.0 $\mu$m outside it) and the cell edge (within 0.8 $\mu$m of the cell boundary) were defined from the cell and nucleus boundaries. The ten cells drawn from each shape share its layout and differ in counts, random fields and molecule draws. The nine compartment modules were the nucleolus, nuclear speckles, nuclear periphery, perinuclear region, ER, mitochondria, Golgi, vesicles and cell edge. The five placement mechanisms were uniform placement within the region; a surface mechanism, Gaussian in distance to the region boundary ($\sigma = 0.10$–$0.20$ $\mu$m); a gradient, the region plus an exponential tail outside it (decay $0.2$–$0.5$ $\mu$m); clustering, a Thomas process with foci in the region (foci $\sigma = 0.08$–$0.20$ $\mu$m; $0.5$–$1.5$ foci per $\mu m^2$); and patchiness, a log-Gaussian Cox field restricted to the region (correlation length $0.2$–$0.5$ $\mu$m). The background genes, ten of each type, were uniform over the cell; clustered in 8–20 whole-cell foci per cell ($\sigma = 0.15$–$0.35$ $\mu$m); or patchy, following a whole-cell log-Gaussian field (correlation length $0.4$–$1.0$ $\mu$m). Molecule counts followed a negative binomial distribution: the mean was log-uniform on 3–100 molecules per cell and scaled by a cell-specific depth factor, and the size parameter was log-uniform on 2–30. Counts and random fields were fixed within a panel across $\lambda$.

**Methods compared.** All methods received molecule identifiers, cell identifiers, gene identifiers and coordinates. InSpace-C scores were $\log_2 C$ over $[0, r)$. InSTAnT[3] (sc-instant 1.2.1) computed proximal pairs at distance $r$ (minimum gene count 20), followed by global colocalization (cell-wise $\alpha = 0.001$, no minimum transcript count, high-precision mode); its score was $-\log_{10}$ of the conditional Poisson binomial $P$ value. Bento[4] scores were colocation quotients at radius $r$, computed with a port of Bento's per-cell quotient, requiring at least ten molecules of each gene in a cell, and averaged on the linear scale over eligible cells. Directed scores were symmetrized by averaging both directions, and pairs that a method did not score were treated as missing.

**Spatial truth and display.** As a gene-level spatial truth, we computed for each gene pair $P(\|X_g - Y_h\| < r) - P(\|U - V\| < r)$ from the generating densities, where $U$ and $V$ are uniformly placed molecules, averaged over cells. For display only (Supplementary Fig. 6a), each method's matrix was background-subtracted on its own scale as $X_{ab} - m_a - m_b + M$, where $m_g$ is gene $g$'s mean score with the 30 background genes and $M$ is the background–background mean; for Bento this used $\log_2$ of the colocation quotient over its within-cell random-label expectation. Each matrix was then shown as within-measure percentiles.

**Supplementary Note 3. Liver fractionation reference**

Mouse liver Xenium Prime data[5] (GSE310392; samples GSM9295277, GSM9295280, GSM9295283 and GSM9295287; 35,235 hepatocyte-lineage cells) were compared with nuclear and cytoplasmic RNA-seq of fractionated mouse liver.[6] Of the 3,477 genes shared by the two datasets (Supplementary Fig. 4c), we kept the 1,110 genes with measurable and differential nuclear and cytoplasmic expression in the Xenium data (compartment expression $\geq 10^{-4}$, nuclear-versus-cytoplasmic test $q \leq 0.25$), following the HiVis gene filter[7,8] (Supplementary Fig. 4d). Genes with a fractionation nuclear-to-cytoplasmic ratio above 2 or below 0.5 formed the 20 nuclear and 967 cytoplasmic reference genes (Supplementary Fig. 4e). Readouts, AUROC and bootstrap intervals were as for the APEX-seq reference (Methods), with 2,000 resamples and the Bento contrast $p_{\text{nuclear}} - p_{\text{cytoplasmic}}$. Coverage is the fraction of cells and of cell–gene pairs for which each method returned an output (Supplementary Fig. 4h).

**Supplementary Note 4. APEX-seq co-occurrence reference and scaling runs**

**Data and method scores.** The HEK293T Xenium data[1,2] comprised 121,343 quality-controlled cells with 275,225,423 transcripts of 290 genes after removal of 61 negative-control features, and all three methods read the same transcript table. InSpace-C scores were $\log_2 C$ over $[0, 2)$ $\mu$m. InSTAnT (`run_ProximalPairs`, distance threshold 2 $\mu$m)[3] scores were the fraction of cells expressing both genes in which the cell-wise proximal-pair test gave $P < 0.001$. Bento (`tl.coloc_quotient`, radius 2 $\mu$m)[4] scores were the mean $\log_2$ colocation quotient, averaged over both directions.

**APEX-seq reference.** The reference was APEX-seq[9] enrichment ($\log_2$ fold change) of 250 panel genes in eight compartments: nucleus, nucleolus, nuclear lamina, nuclear pore, ER membrane, ER lumen, outer mitochondrial membrane and cytosol. Each gene's profile was row-centred and L2-normalized, and the reference for a gene pair was the cosine of the two profiles. Each gene was assigned to its compartment of maximal enrichment, and a pair was classified as same compartment, adjacent compartment or other, with adjacency defined a priori for nucleus–nucleolus, nucleus–lamina, nucleus–pore, lamina–pore, pore–cytosol, pore–ER membrane, cytosol–ER membrane, cytosol–outer mitochondrial membrane and ER membrane–ER lumen. Top-ranked pairs were taken with ties broken by pair identifier. In the matrices of Fig. 2d, genes were ordered by APEX-seq compartment and then by decreasing enrichment, without using any method score, and each measure was shown as within-measure percentiles.

**Permutation test and lowest-ranked pairs.** In a secondary analysis, each method was ranked within all pairs it scored, and the compartment relation of its top 1%, 5% and 10% pairs was compared with 5,000 gene-label permutations that shuffle APEX-seq profiles and compartment assignments across genes while keeping method scores fixed. The bottom 1%, 5% and 10% pairs of each method, ranked by ascending score on the same 13,299 shared pairs with ties broken by pair identifier, were classified in the same way and compared with the composition of all shared pairs (Supplementary Fig. 6c).

**Scaling runs.** Nested subsets of 200, 1,000, 5,000, 20,000 and 60,000 cells, and all cells, were taken as prefixes of one seeded random ordering. Each method and subset ran as a separate job on a dedicated node with 48 CPU cores and 384 GiB of memory, using 48 threads. Run time was the wall-clock time of the method call, excluding input loading and output writing, and peak memory was the peak resident set size of the method process (Fig. 2e and Supplementary Fig. 6d,e).

**Supplementary Note 5. B and plasma cell analyses**

**Cohort.** Of 68 tumour sources, nine breast cancer sources on a 280-gene panel with fewer than three plasma-cell markers, the minimum for cellAdmix marker mode, were excluded, and one ovarian cancer source with 8 B cells after filtering entered no analysis. Breast cancer sources include triple-negative breast cancer (TNBC) and lung cancer sources include non-small-cell lung cancer. Sources are not claimed to be independent donors.

**Removal of molecules from neighbouring cells.** For each source, cellAdmix[10] (v0.1.0) learned molecular neighbourhood patterns by non-negative matrix factorization (ten factors, 20 nearest molecules within the cell, 10,000 neighbourhoods sampled across cell types), and a conditional random field assigned each molecule of B and plasma cells to a factor. Factors were attributed to lineages by enrichment of lineage markers (package default, FDR $< 0.1$), using one marker table per cancer type that listed canonical markers of immune, stromal and parenchymal lineages from published atlases.[11–21] A factor flagged as foreign was kept in a cell type when its median per-cell fraction there was at least 0.1, the threshold of the package function `check_f_rm`. Molecules of the remaining foreign factors were removed from B and plasma cells.

**Differential radial localization.** Within a source, cell–gene pairs with at least three transcripts were used, and a gene entered the source when at least 20 such cells were present in each state. The null distribution of $T$ enumerated all $2^{k-1}$ sign patterns of the source contrasts when $k \leq 15$ and otherwise drew 19,999. $W_2$ is the exact 2-Wasserstein distance, signed by the mean shift. Log-normalized counts are $\log(1 + x)$ of counts scaled to $10^4$ per cell, and the abundance difference was averaged over sources with at least 20 cells of each state. **Differential co-occurrence and neighbour controls.** Ordered pairs of molecules within 2 $\mu$m were counted in each cell and summed over cells to give $C_{ab} = J_{ab}T/(M_a M_b)$, where $J_{ab}$ is the number of pairs with labels $a$ and $b$, $M_a$ and $M_b$ the numbers of pairs with $a$ at the first and $b$ at the second molecule, and $T$ the total number of pairs. A pair was supported in a source and state when the state had at least 50 cells containing RNA, the expected joint count $M_a M_b/T$ was at least 100, at least 5% of cells contained both genes and $J_{ab} > 0$. Of the four MERFISH sources, the colorectal source supported 92 of the 5,414 tested pairs, and the three breast sources, whose panels carry no HLA class II genes, one pair in total. The permutations of the B and plasma labels were shared by all pairs, with support fixed at the observed labels. For the neighbour controls, the MHC-II count of each cell was the sum of its HLA class II transcripts (*HLA-DR*, *-DP*, *-DQ*, *-DM* and *-DO* genes on the panel) before filtering, and a cell whose centroid lay within the radius was an MHC-II-high neighbour when this count was at least one and at least the median of the plasma cells in the same source. The 90th percentile and a single transcript were alternative thresholds at 15 $\mu$m. Cells without counts (1–3% of cells in four sources) were not MHC-II-high. B-lineage neighbours were B and plasma cells. The controls used the 18 sources whose panels included *HLA-DRA* for the radial test and 13 of the 16 sources for the permutation test, the other three having no HLA class II genes on the panel.

**Co-occurrence programs and TNBC replication.** Genes marking other lineages were removed, and genes with at least ten supported pairs were kept. Masked entries of $A$ were excluded from the fit rather than set to zero. Genes measured on different platforms share no cells and can share a program only through genes measured on both. The symmetric NMF used a ridge penalty ($\lambda = 0.3$) and damped multiplicative updates. The number of programs was the smallest $k$ whose held-out error, with 10% of observed pairs hidden in five splits, lay within one standard error of the minimum. Random starts were aligned by Hungarian assignment,[22] and a gene was a stable member of a program when it carried at least 30% of its weight on that program in at least 80% of starts. Programs were numbered by anchor genes. ER-targeted proteins were UniProtKB/Swiss-Prot[23] (release 2026_03) entries with a signal peptide or with a transmembrane segment that is not a C-terminal tail anchor, excluding mitochondrial and peroxisome-only entries. In the TNBC replication, pairs supported in at least three of the six TNBC sources were testable, with the median $\log_2 C$ over them as value, and the genes considered were those of the discovery fit with at least ten testable partners (118 in B cells, 240 in plasma cells). Each gene's TNBC weights were fitted with the discovery weights of all other genes held fixed, so its own discovery weight was not used, and the permutations moved the TNBC gene labels with the discovery weights fixed. The stable members of B3, P4 and P5 were placed mostly on neighbouring programs (B2, P2 and P3). As supportive readouts (Supplementary Table 2), the mean $\log_2 C$ among the stable members of each program was compared with random gene sets matched on testable pairs per gene, and programs were also fitted on the TNBC sources alone and matched to the discovery programs by Hungarian assignment.

**Consistency across sources, cancer types and platforms.** For each source, we recorded the fraction of detected genes whose shift in that source had the cohort direction. The radial test was rerun with identical settings without the sources of each cancer type, of each platform or of the TNBC lymph nodes (17 reruns), and with only breast, lung or colorectal cancer, Xenium, or MERSCOPE and MERFISH sources (5 reruns), the groups with at least six sources (Supplementary Table 2). Many genes reached the minimum of six sources only with the TNBC lymph-node sources, without which 98 genes could be tested. The permutation test was rerun without each of the seven cancer types and with only MERSCOPE, MERFISH, breast or colorectal sources (Supplementary Table 2). In each rerun, every gene or pair significant in the cohort was classed as significant in the cohort direction, same direction but not significant, not tested, or reversed. Program support in a source was the mean $\log_2 C$ among stable members minus that among all supported pairs. Programs were refitted with the same $k$ without each of the nine cancer types and with only MERSCOPE and MERFISH, Xenium Prime 5K, lung or breast sources, matched to the full programs as above and assessed when at least five members and ten testable member pairs remained (Supplementary Table 2). No detected gene reversed direction in any of the 22 radial reruns, none of the 2,127 differing pairs reversed with $q < 0.05$, and without any single cancer type B1–B3 and P1 kept a weight correlation of at least 0.92, P5 of at least 0.87, and P2 and P3 of at least 0.74 and 0.63, whereas B5 fell to $-0.26$ without the colorectal sources, P4 was lost without the colorectal, liver or melanoma sources, and B4 ranged from $-0.12$ to 1.00.

**Expression gene-set enrichment.** Differential expression between plasma and B cells was tested with sources as replicates, because tests that treat cells as independent replicates inflate false discoveries.[24] For each gene, the per-source difference in mean log-normalized expression (plasma − B; sources with at least 20 cells of each state) was tested against zero with a one-sample $t$-test over the sources measuring the gene, for the 5,371 genes measured in at least six sources, and $P$ values were adjusted by the Benjamini–Hochberg procedure.[25] Genes were ranked by signed $-\log_{10} P$[26] and tested against the MSigDB Hallmark collection[27] by preranked gene-set enrichment analysis[28] in GSEApy[29] (set size 5–500, 1,000 permutations), with the tested genes as background.[30] $q$ values were computed over the 50 sets from the nominal $P$, with

$P = 0$ set to $1/1{,}001$ (Supplementary Fig. 10e). As a check that does not require genes to be matched across panels,[31] the enrichment was also run in each source, with genes ranked by that source's difference and that source's genes as background, and the direction of each set was compared across sources: the unfolded protein response was higher in plasma cells in all 47 sources in which it could be tested, mTORC1 signalling in 42 of 50 and the Allograft Rejection set higher in B cells in 49 of 58.

**Interpretation of the enriched sets.** The sets match known programmes of plasma-cell differentiation. XBP1 and the unfolded protein response expand the secretory apparatus,[32–35] and the leading edge here contained all three branches (*XBP1*, *ERN1*, *EIF2AK3*, *ATF6*, *ATF4*). mTORC1 is required for plasma-cell generation and antibody synthesis and induces a UPR-related programme before secretion begins;[36–38] its leading edge contained amino-acid transporters (*SLC7A5*, *SLC1A5*) and ER genes. The Allograft Rejection set was carried by antigen-presentation and B-cell activation genes (*HLA-DRA*, *HLA-DMA*, *CD40*, *CD80*, *CD86*, *PTPRC*, *LYN*, *IRF8*), in line with the extinction of the mature B-cell programme and of CIITA-dependent MHC-II by Blimp-1.[39–41] Fatty-acid metabolism is consistent with XBP1-driven phospholipid synthesis for ER expansion and with the rise in oxidative metabolism during plasma-cell differentiation,[42–44] although a coordinated increase in fatty-acid oxidation enzymes in plasma cells has not been described. The UV Response Up set has no specific counterpart in plasma-cell biology; its leading edge consisted of stress and apoptosis genes (*ATF3*, *CASP3*, *BAK1*, *BCL2L11*).

**Epithelial–mesenchymal transition gene set.** The Hallmark epithelial–mesenchymal transition (EMT) set was enriched in plasma cells (NES 1.73, BH $q = 0.012$). The enrichment was carried by the stromal and extracellular-matrix (ECM) genes of the set, such as *COL4A1*, *COL5A1*, *FN1*, *CTHRC1* and *TNC* (NES 2.07, nominal $P < 0.001$; genes that are fibroblast, mural or endothelial markers in the cellAdmix marker tables or belong to collagen and ECM structural families, 72 tested). The remaining EMT genes were not enriched (NES 1.09, nominal $P = 0.31$). Each ECM gene differed little between the two states: the mean difference in log-normalized expression was mostly below 0.3, against 1–4 for B-cell and plasma-cell markers. In both plasma and B cells these transcripts lay nearer the cell boundary than other transcripts. Their median radial position was 0.10 above that of the median gene in the same source, in 92% of 105 gene–source pairs, as expected for molecules assigned to a cell from outside it. In imaging-based spatial transcriptomics, transcripts are misassigned between neighbouring cells through segmentation errors and lateral diffusion,[10,45] and through cells that overlap along the $z$ axis of the section.[46] This contamination scales with the local abundance of the source cell type and persists in cells classified as singlets,[45] and reduced molecular specificity of this kind can distort differential expression.[47] Three corrections reduced the ECM differences but did not remove them: molecule filtering with cellAdmix, exclusion of cells with a stromal cell within 15 $\mu$m, and restriction of expression to the cell interior (radial position $< 0.8$). The EMT enrichment remained in each case (NES 1.62–2.37). In tumours, plasma cells disseminate into the tumour bed along fibroblastic tracks, whereas B cells concentrate in tertiary lymphoid structures.[48] The most likely source of the EMT signal is therefore the denser stromal background around plasma cells, and we do not interpret it as an epithelial–mesenchymal programme in plasma cells.

### Supplementary Note 6. Cytos datasets

RNA–RNA co-occurrence over $[0, 2)$ $\mu$m was computed with all counted genes as the background in A549 cells (MBA-0832 release) from vehicle-treated wells carrying non-targeting guides[49] and in HeLa cells from DMSO-treated wells[50], pooling counts over wells within each dataset. The matrices show a fixed set of 74 genes: 9 mitochondrially encoded genes, 33 ribosomal-protein genes and 32 protein-coding genes that were abundant in both datasets, chosen by abundance alone (Supplementary Fig. 8a,b).

**Supplementary Note 7. RNA–DNA co-occurrence**

DNA loci were imaged at 25-kb resolution and assigned to 100-kb genomic bins from an annotation independent of the RNA labels. The rank of a gene's cognate bin was one plus the number of candidate bins with a strictly higher score, and the top 1% comprised the $\lceil 0.01 \times \text{candidates} \rceil$ best-ranked bins. Ties for the peak were resolved towards the transcription start site (TSS), and the peak was counted as recovered when its midpoint lay within 1 Mb of the TSS. The chromosome 11 heatmap (Fig. 2h) shows the 120 genes of the 8,865-gene set on that chromosome with the most same-chromosome intron–DNA pairs, ordered by TSS, across 100-kb bins; chromosome 11 has no bins without probes. The heatmaps of Supplementary Fig. 9d show all genes of the set on the two chromosomes with the highest (16 and 15) and the two with the lowest (18 and 19) top-1% recovery, excluding chromosome 11 and the X chromosome. The example nucleus (Supplementary Fig. 9a) is the nucleus with the median number of intron spots among those with intron and DNA spots in the first field of view of replicate 1. The example gene is the one among the 120 genes of Fig. 2h with the most intron spots that all have DNA of the gene's own bin within 0.5 $\mu$m, with ties broken by the number of such loci.

49. Element Biosciences: Cytos multimodal pooled screening in A549 cells: MBA-0832. Element Biosciences public dataset Dataset; analysed release MBA-0832, vehicle wells with non-targeting guides; accessed 21 October 2026.

50. Element Biosciences: Profiling Compound Response of Tunicamycin with Cytos. Element Biosciences public dataset Dataset; Cytos-3Prime_HeLa-Tunicamycin, Run2_WTA_3; DMSO wells; accessed 21 October 2026.

51. Xia, C., Fan, J., Emanuel, G., Hao, J., Zhuang, X.: Spatial transcriptome profiling by MERFISH reveals subcellular RNA compartmentalization and cell cycle-dependent gene expression. Proc Natl Acad Sci U S A **116**(39) (2019) 19490–19499

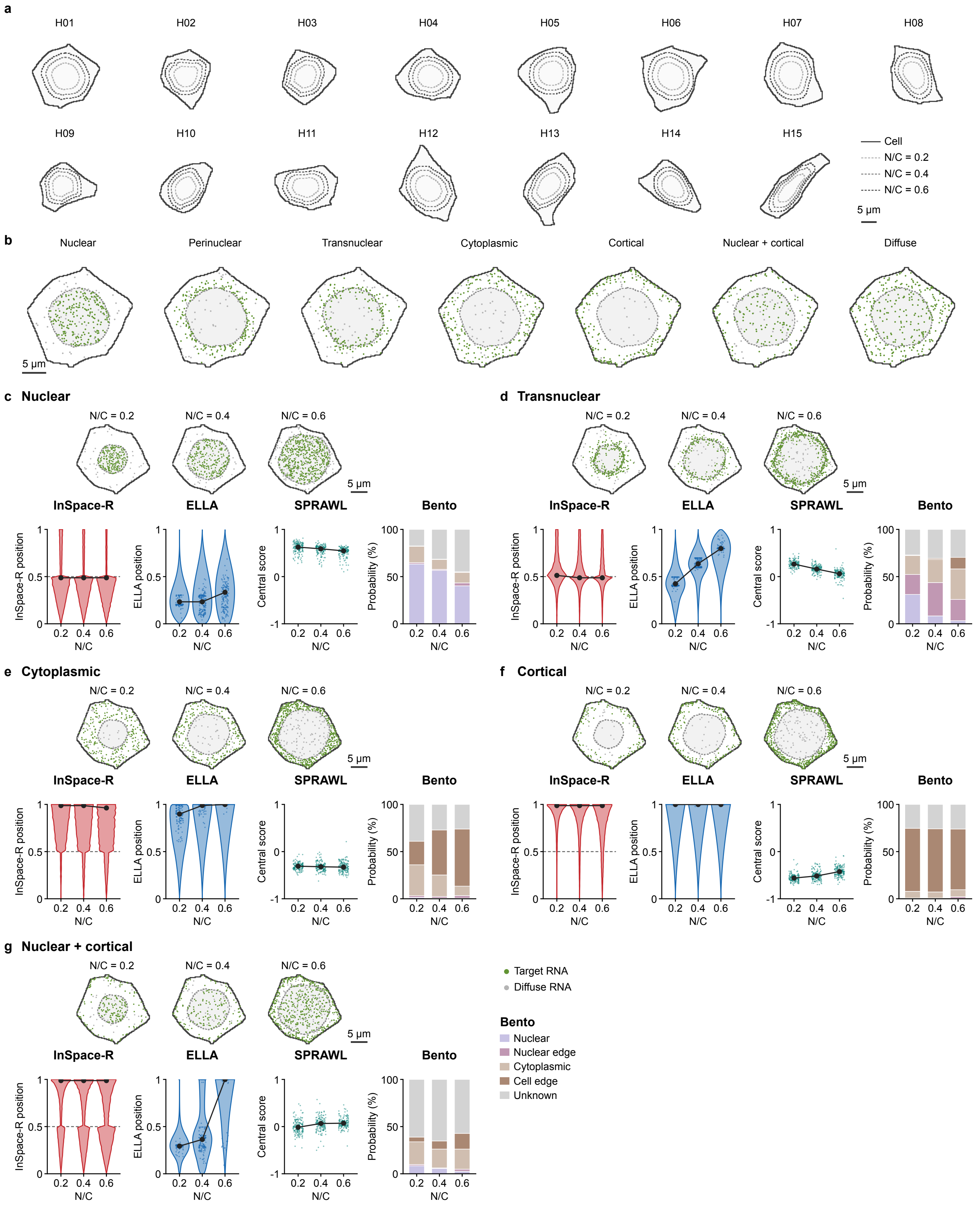


**Supplementary Figure 1 Simulation templates, planted patterns and native readouts across N/C. a**, The 15 cell templates, cells H01–H15 of the HEK293T Xenium data, each with its nucleus at a nucleus-to-cell area ratio (N/C) of 0.2, 0.4 and 0.6. The cell outline is fixed and only the nucleus is enlarged. **b**, The seven planted patterns in template H01 at N/C = 0.4. Each panel shows all molecules of the ten genes assigned to that pattern. **c–g**, The nuclear, transnuclear, cytoplasmic, cortical and nuclear + cortical patterns, shown as for the perinuclear pattern in Fig. 1c, with template H01 at N/C = 0.2, 0.4 and 0.6 (top) and the native readouts of InSpace-R, ELLA, SPRAWL and Bento (bottom).

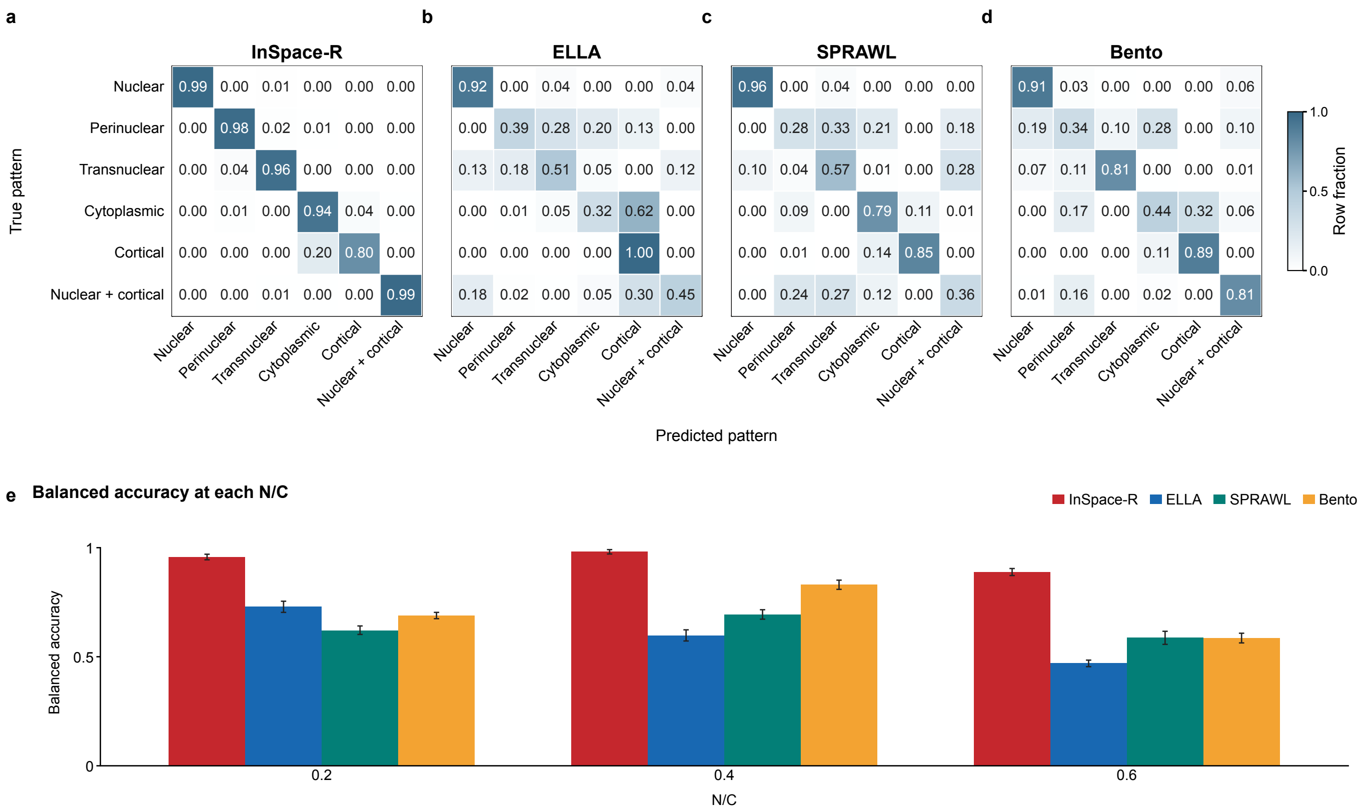


**Supplementary Figure 2 Six-pattern recognition. a**–**d**, Confusion matrices of the six-pattern recognition in Fig. 1d for InSpace-R (**a**), ELLA (**b**), SPRAWL (**c**) and Bento (**d**). **e**, Balanced accuracy of each method at each N/C. Error bars show 95% CIs.

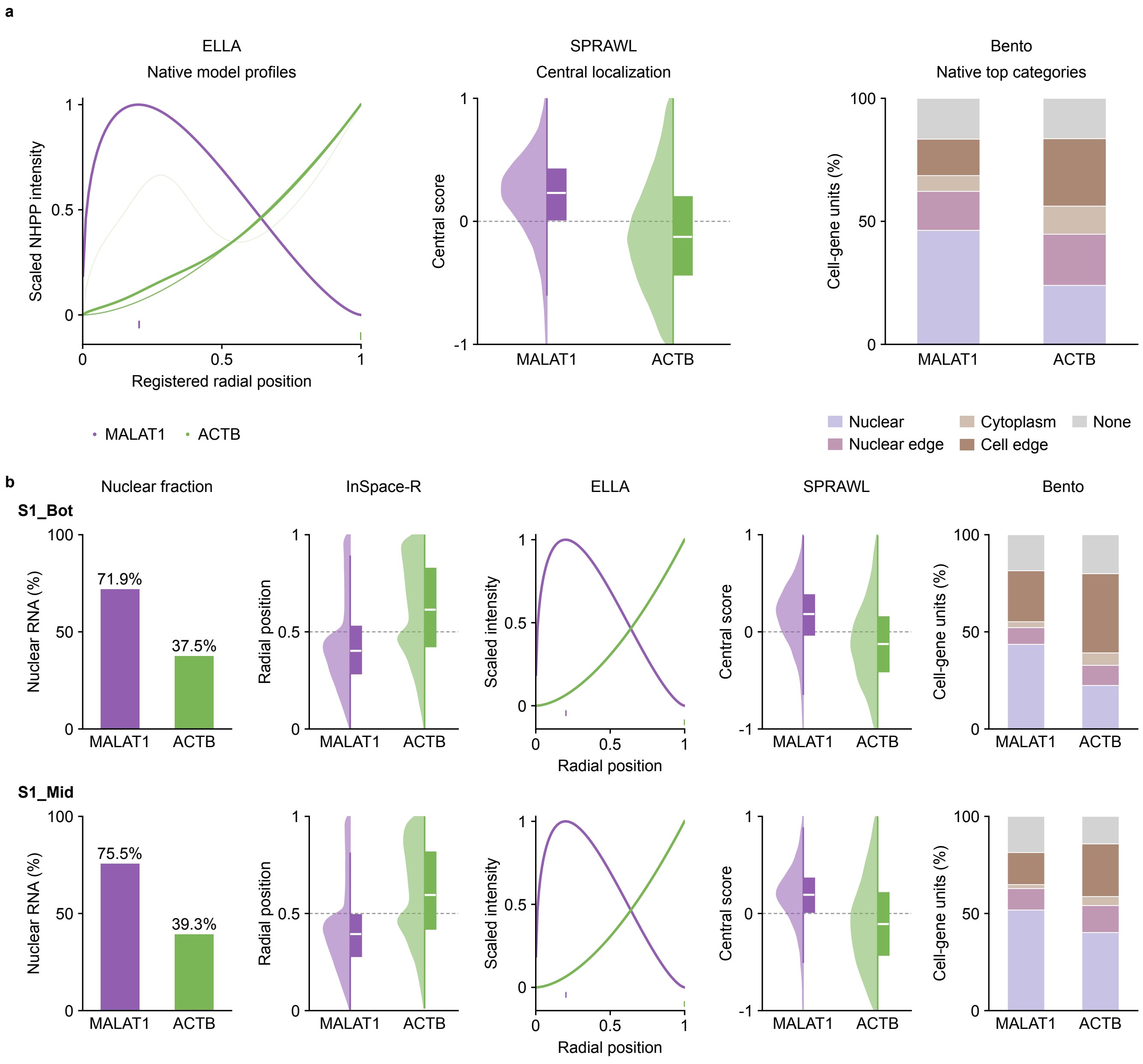


**Supplementary Figure 3** (1/3, continued on the next page)

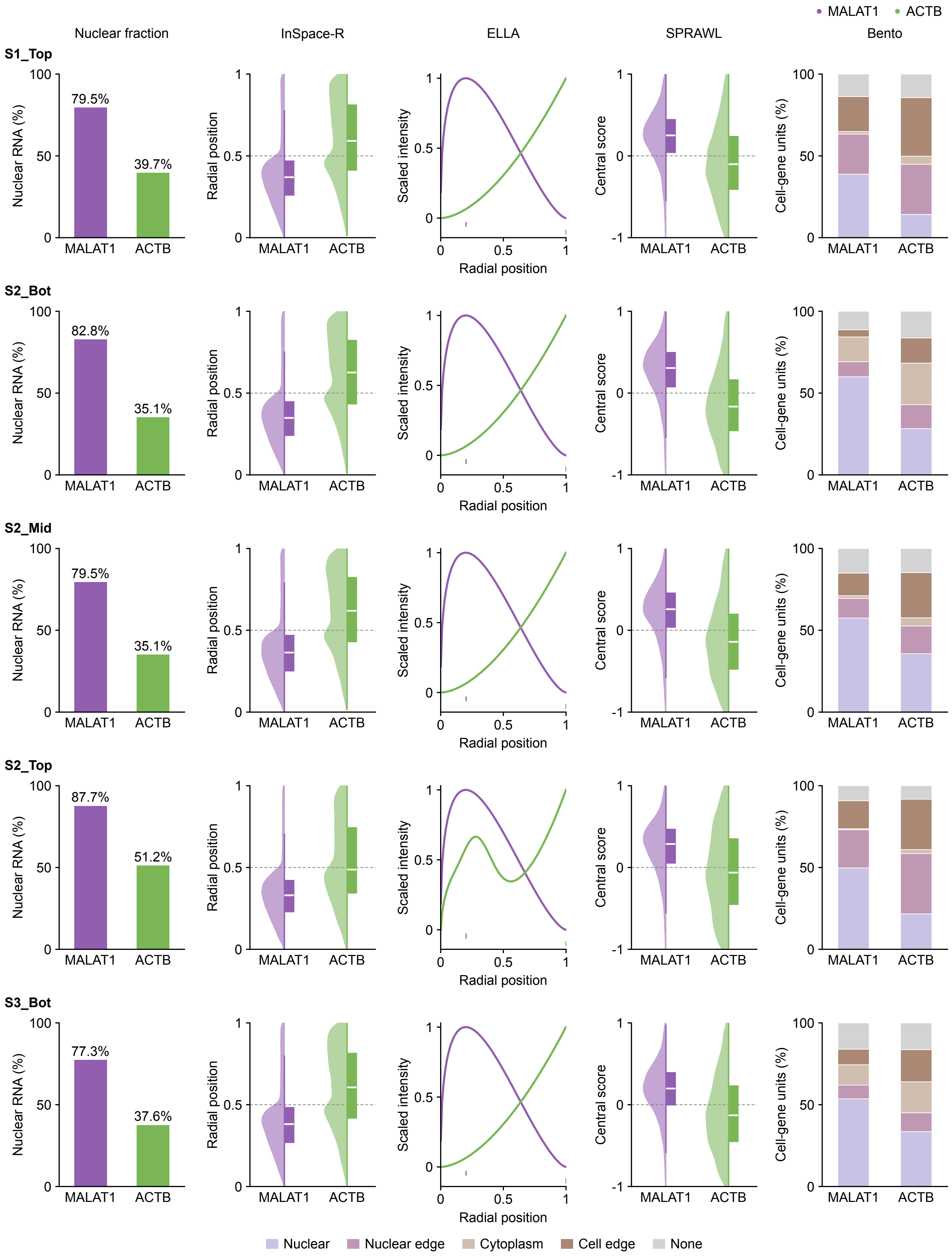


**Supplementary Figure 3** (2/3, continued on the next page)

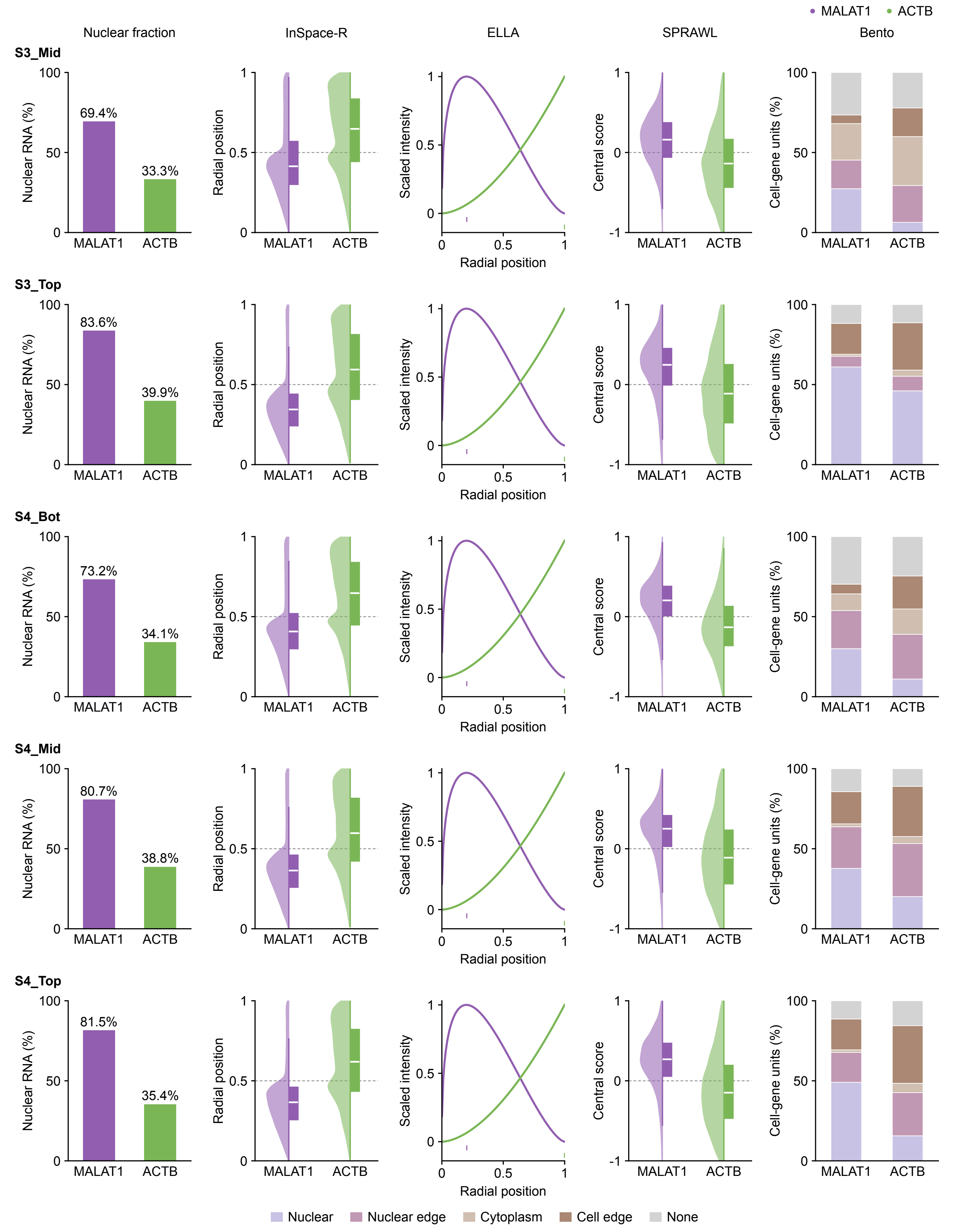


**Supplementary Figure 3** (3/3) **MALAT1 and ACTB pooled and in each of the 12 datasets. a**, Native readouts of ELLA, SPRAWL and Bento for MALAT1 and ACTB, pooled over the 12 Xenium datasets in Fig. 1e. Thin ELLA lines show individual datasets and thick lines their mean. **b**, The 12 datasets, two on the first page and five on each following page. Columns show the nuclear fraction and the readouts of InSpace-R, ELLA, SPRAWL and Bento, on the same axes in every row.

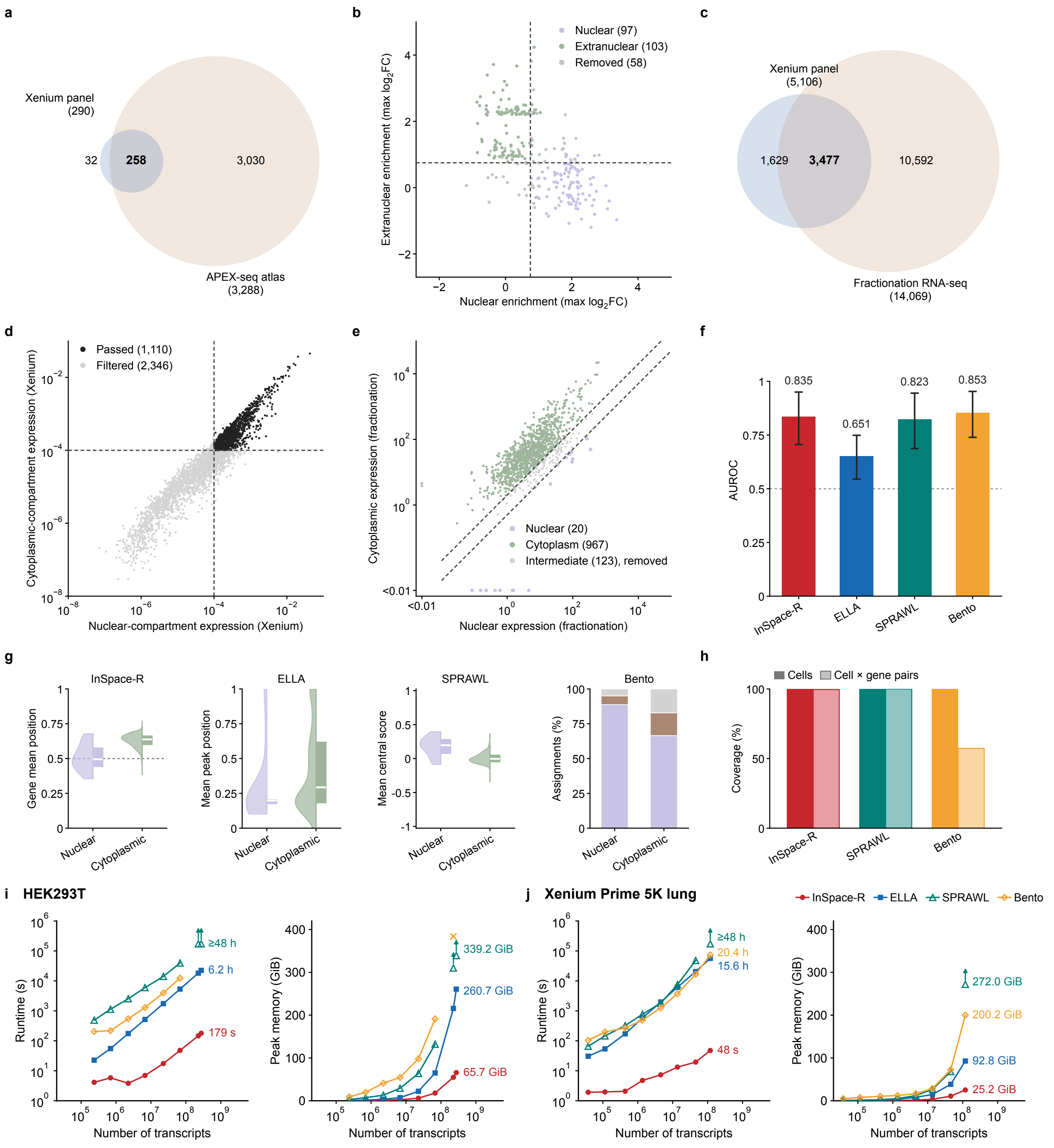


**Supplementary Figure 4 HEK293T APEX-seq and mouse liver fractionation references, and runtime and memory.** **a**, Overlap between the genes on the HEK293T Xenium panel and the genes in the APEX-seq atlas. **b**, Largest published $\log_2$ fold change in the nuclear and in the extranuclear compartments for each shared gene, coloured by whether the gene was kept as a nuclear or extranuclear reference gene or removed. Dashed lines mark the fold-change cutoff of 0.75. **c**, Overlap between the genes on the mouse liver Xenium panel and the genes detected by fractionation RNA-seq. **d**, Nuclear and cytoplasmic compartment expression of the shared genes in the Xenium data. **e**, Nuclear and cytoplasmic expression of the passing genes in the fractionation data, which defines the nuclear and cytoplasmic reference genes. Dashed lines in d and e mark the selection thresholds. **f**, AUROC of each method against the liver reference. Error bars show 95% CIs. **g**, Per-gene readouts of each method for the nuclear and cytoplasmic reference genes. **h**, Fraction of cells and of cell $\times$ gene pairs with output from InSpace-R, SPRAWL and Bento. **i**,**j**, Runtime (left) and peak memory (right) of the four methods at increasing numbers of transcripts in HEK293T (**i**) and Xenium Prime 5K lung (**j**) data. Upward arrows mark runs stopped at the 48-h limit, and the cross marks a run that ran out of memory.

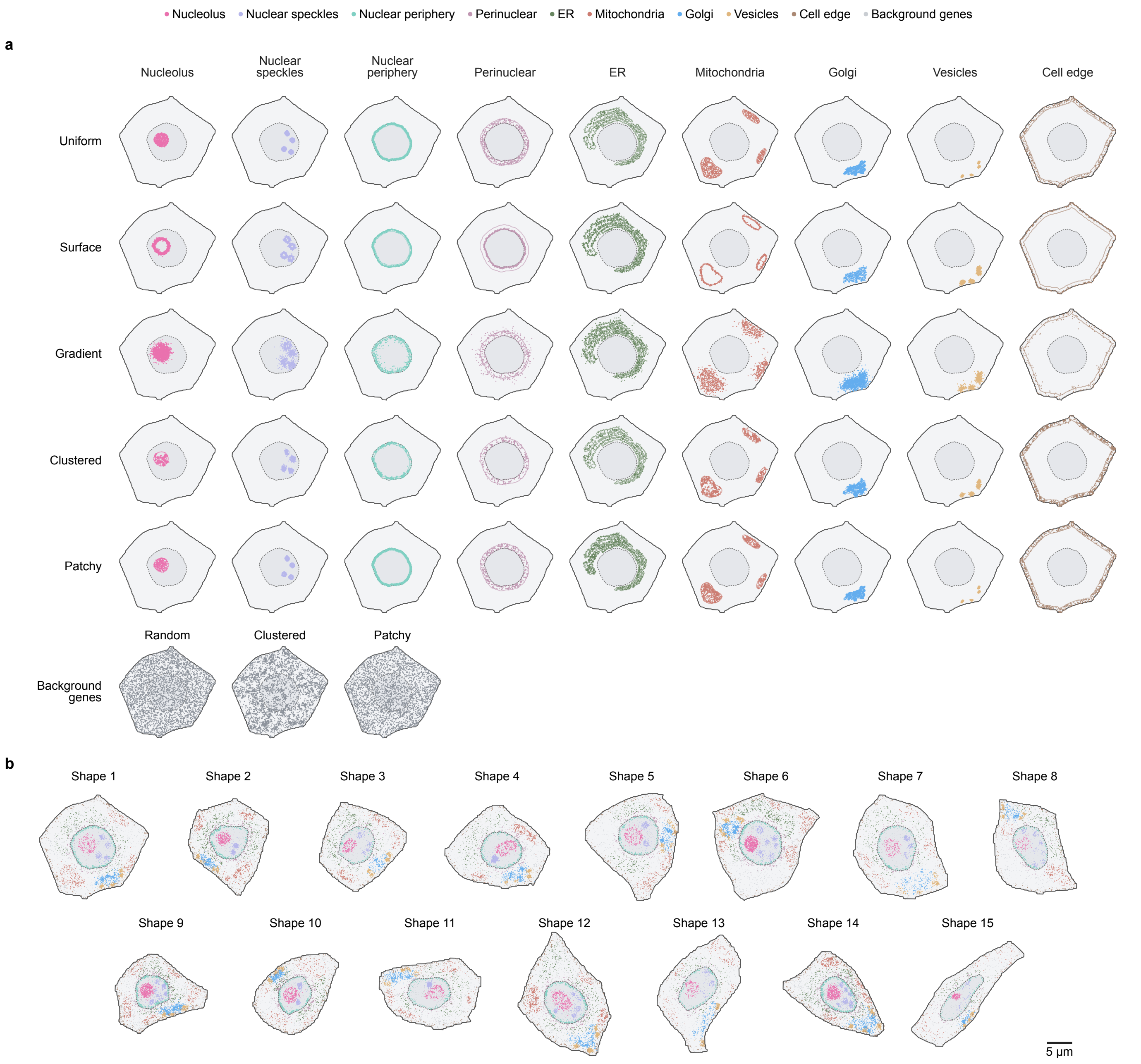


**Supplementary Figure 5 Design of the simulated RNA co-occurrence data. a**, Molecules of the nine compartment modules under five placement mechanisms, and of the three background-gene types; lines, module regions. **b**, The 15 simulated cell shapes, taken from HEK293T cells at a nucleus-to-cell area ratio of 0.2.

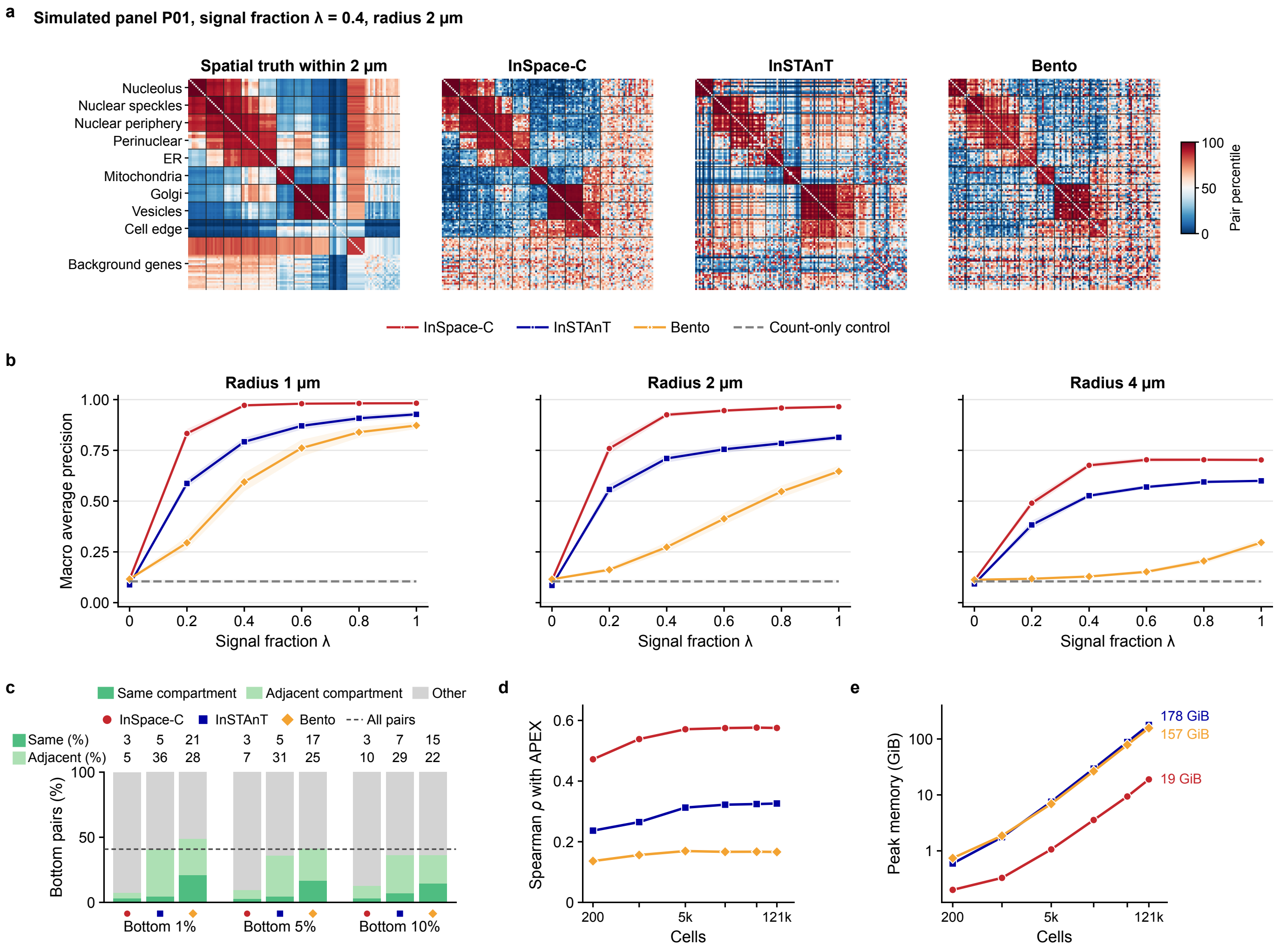


**Supplementary Figure 6 Simulation results across settings and further HEK293T APEX-seq readouts.** **a**, Background-subtracted gene-pair scores in one simulated panel ($\lambda$ = 0.4, 2 $\mu$m), shown for the spatial truth and each method as within-measure percentiles. **b**, Macro average precision as a function of $\lambda$ at radii of 1, 2 and 4 $\mu$m. Lines show the mean and bands the s.d. ($n = 10$ simulated panels). **c**, APEX-seq compartment relation of each method's bottom 1%, 5% and 10% gene pairs in HEK293T, the pairs it scores as most depleted. Numbers above the bars give the percentages of same- and adjacent-compartment pairs, and the dashed line marks the share of same- or adjacent-compartment pairs among all pairs. **d**, Spearman $\rho$ between each method's scores and the APEX-seq cosine as a function of cell number. **e**, Peak memory as a function of cell number.

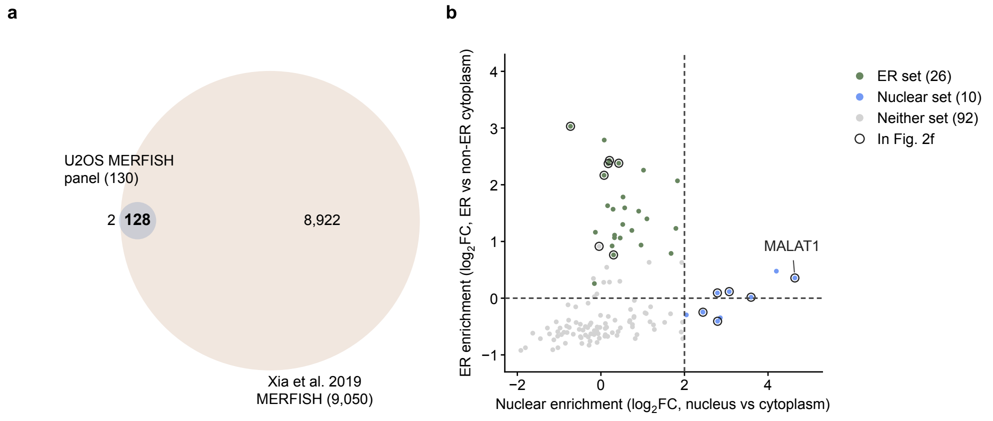


**Supplementary Figure 7 Localization sets of the U2OS panel genes from Xia et al. (2019). a**, Overlap between the genes on the U2OS MERFISH panel and the 9,050 genes analysed by Xia et al.[51] **b**, Nuclear enrichment (nucleus versus cytoplasm) and ER enrichment (ER versus non-ER cytoplasm) from Xia et al. for each shared gene, coloured by the localization set used in Fig. 2f. All thresholds are those of Xia et al.: the ER set comprises genes with an ER $\log_2$ fold change above 0 and the nuclear set genes with a nuclear $\log_2$ fold change above 2, both at a Bonferroni-adjusted $P$ below $10^{-10}$ (dashed lines). These criteria reproduce the published sets of 1,006 ER-enriched and 1,488 nuclear-enriched genes, and the 30 and 17 blank barcodes that Xia et al. reported as ER- and nuclear-enriched. *MALAT1* is the only shared gene in both sets and is assigned to the nuclear set, for three reasons given by Xia et al.: their ER test compares only cytoplasmic molecules, their nuclear calls held after about 1 $\mu$m of the nuclear rim was eroded to exclude perinuclear ER (1,484 of 1,488 genes), and they describe *MALAT1* as a nuclear-enriched lncRNA. Ringed points are the genes in Fig. 2f.

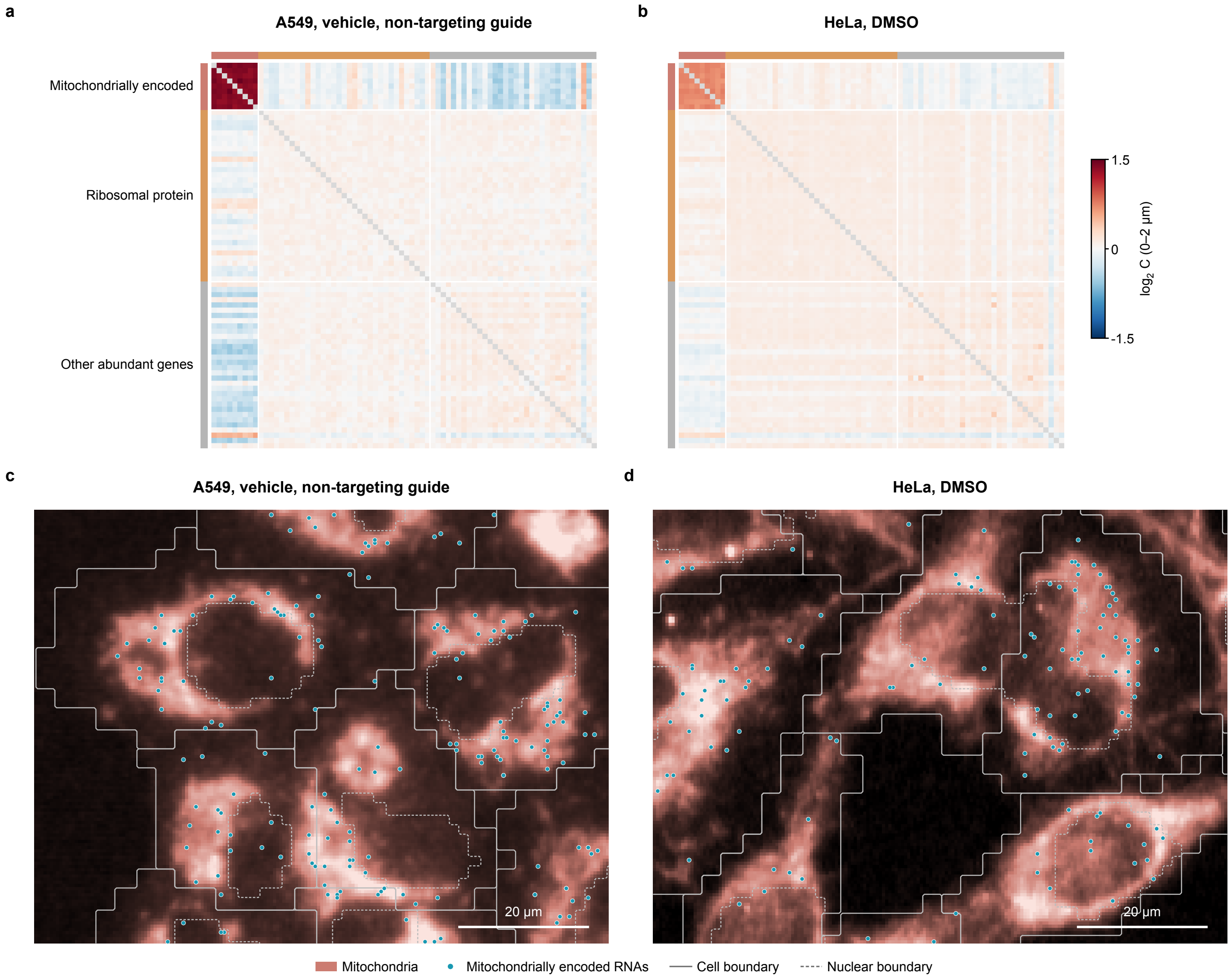


**Supplementary Figure 8 Mitochondrially encoded RNAs co-occur on mitochondria in two Cytos datasets.** **a**,**b**, RNA–RNA co-occurrence ($\log_2$ C, 0–2 $\mu$m) among mitochondrially encoded, ribosomal protein and other abundant genes in A549 (**a**) and HeLa (**b**). **c**,**d**, Fields of control cells showing mitochondria and mitochondrially encoded RNAs in A549 (**c**) and HeLa (**d**).

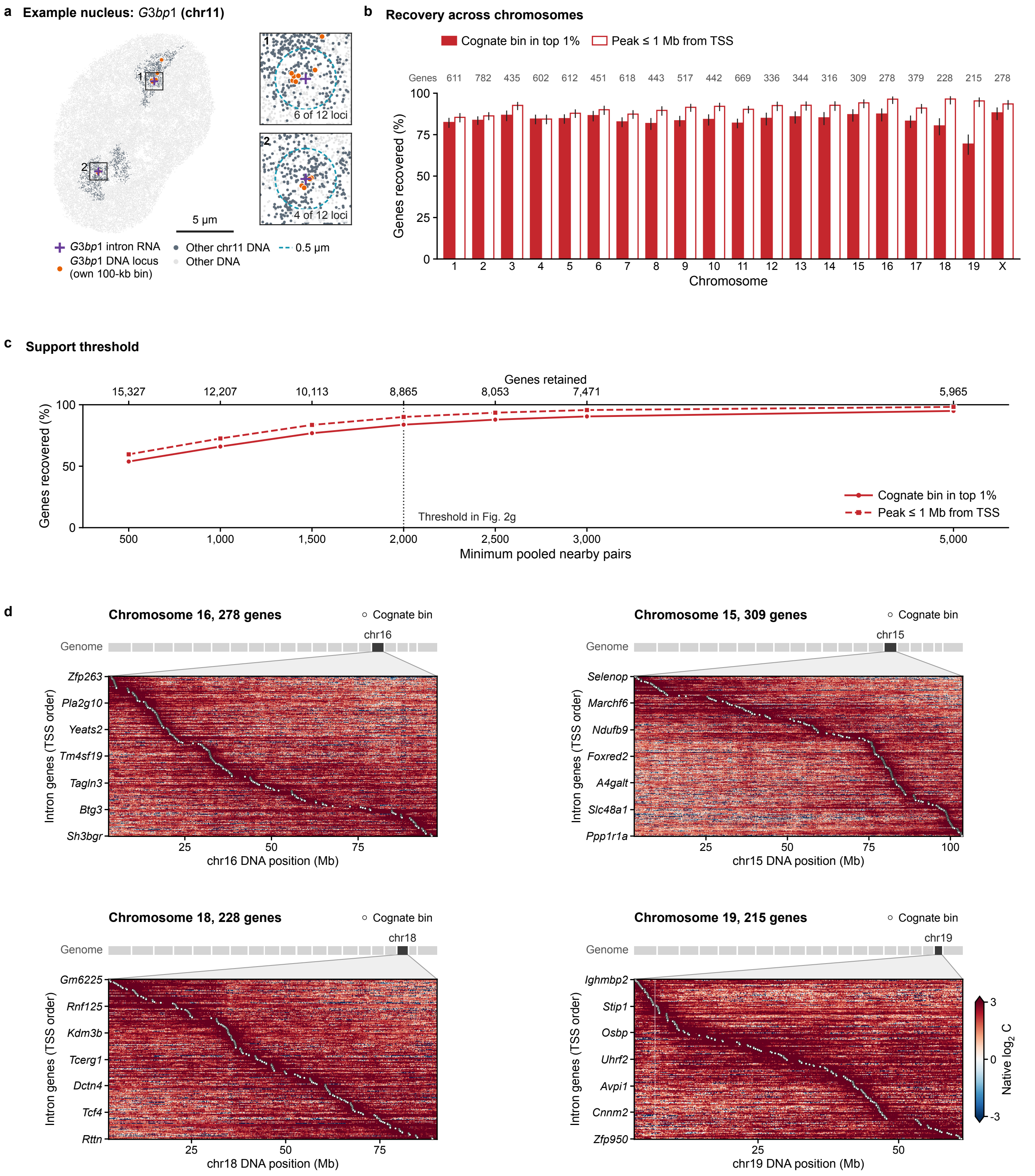


**Supplementary Figure 9 RNA–DNA co-occurrence in mouse embryonic stem cells.** **a**, An mESC nucleus imaged by DNA seqFISH+, with the DNA loci of chromosome 11 in dark grey and those of other chromosomes in light grey. The two intron RNA spots of *G3bp1* (purple crosses), one of the chr11 genes in Fig. 2h, lie in the two chromosome 11 territories, each next to DNA loci of the gene's own 100-kb bin (orange). Enlarged views show each spot with its 0.5-$\mu$m neighbourhood. **b**, Recovery of the source neighbourhood of intron RNAs on each chromosome: the percentage of genes whose cognate DNA bin ranks in the top 1% (filled) or whose peak lies within 1 Mb of the transcription start site (open), with genes per chromosome on top. Error bars show Wilson 95% confidence intervals. **c**, Recovery at each support threshold, with the number of genes retained on the top axis. **d**, Intron RNA–DNA co-occurrence, as in Fig. 2h, for all genes on the two chromosomes with the highest (chr16, chr15) and the two with the lowest (chr18, chr19) recovery in **b**.

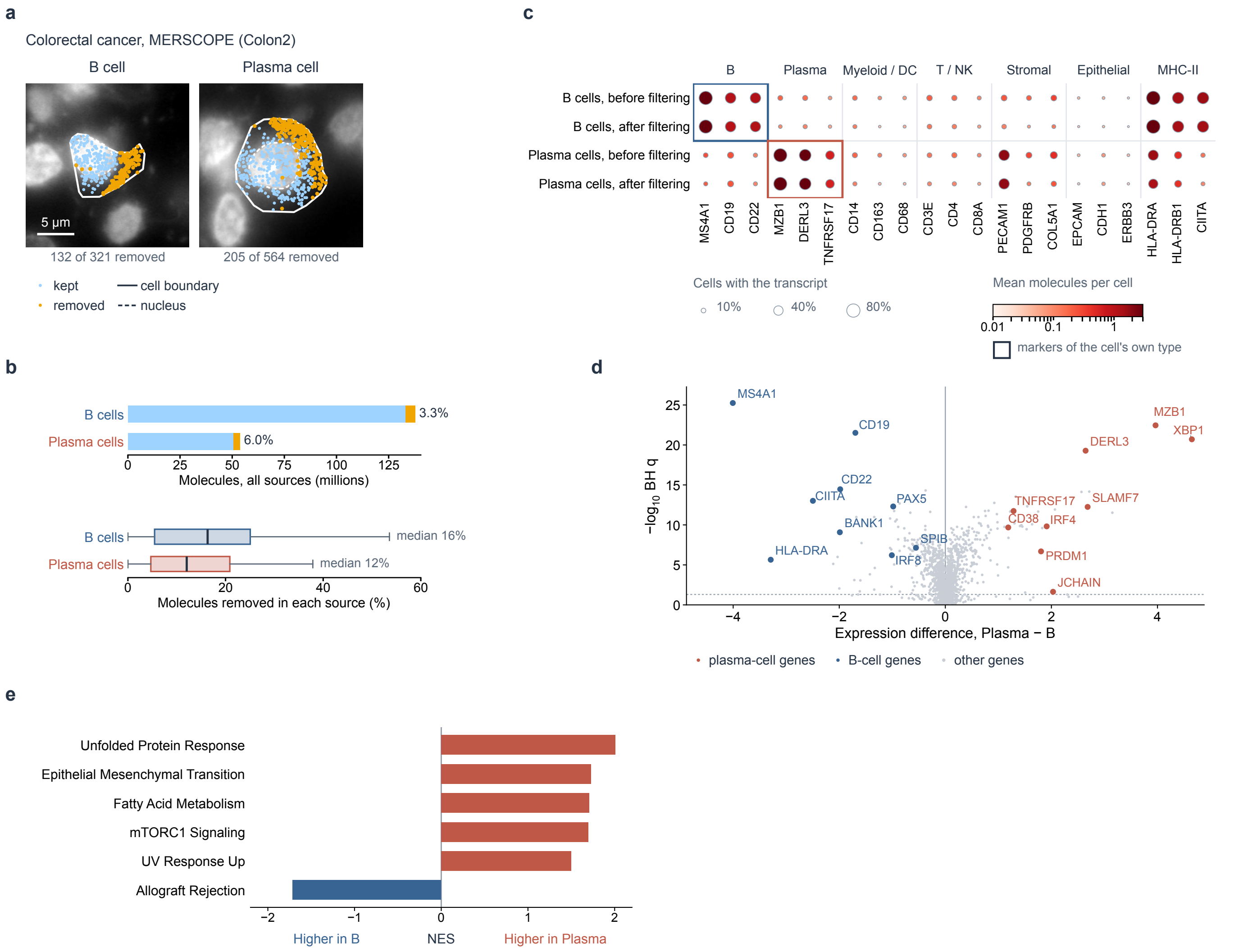


**Supplementary Figure 10 The analysed cells are B and plasma cells, and their molecules are their own.** **a**, A B cell and a plasma cell of the colorectal cancer MERSCOPE source Colon2 on DAPI, with every molecule of the cell, kept or removed by filtering. Solid line, cell boundary. Dashed line, nucleus. Numbers, molecules removed of all molecules of the cell. **b**, Molecules removed by filtering in B and plasma cells, summed over all sources (top; kept and removed, millions of molecules) and in each source (bottom; percentage removed). Boxes, interquartile range. Line, median. Whiskers, 1.5 times the interquartile range. **c**, Lineage markers and MHC-II genes in B and plasma cells before and after filtering. Dot size, fraction of cells with at least one molecule of the gene. Colour, mean molecules per cell. Both are means over the sources that measure the gene. Boxes, markers of the cell's own type. **d**, Difference in mean log-normalized expression between plasma and B cells and its significance over sources for each gene measured in at least six sources. Labelled, canonical plasma-cell (red) and B-cell (blue) genes. Dashed line, $q = 0.05$. **e**, Normalized enrichment score (NES) of each Hallmark gene set with $q < 0.05$. Red, higher in plasma cells. Blue, higher in B cells.

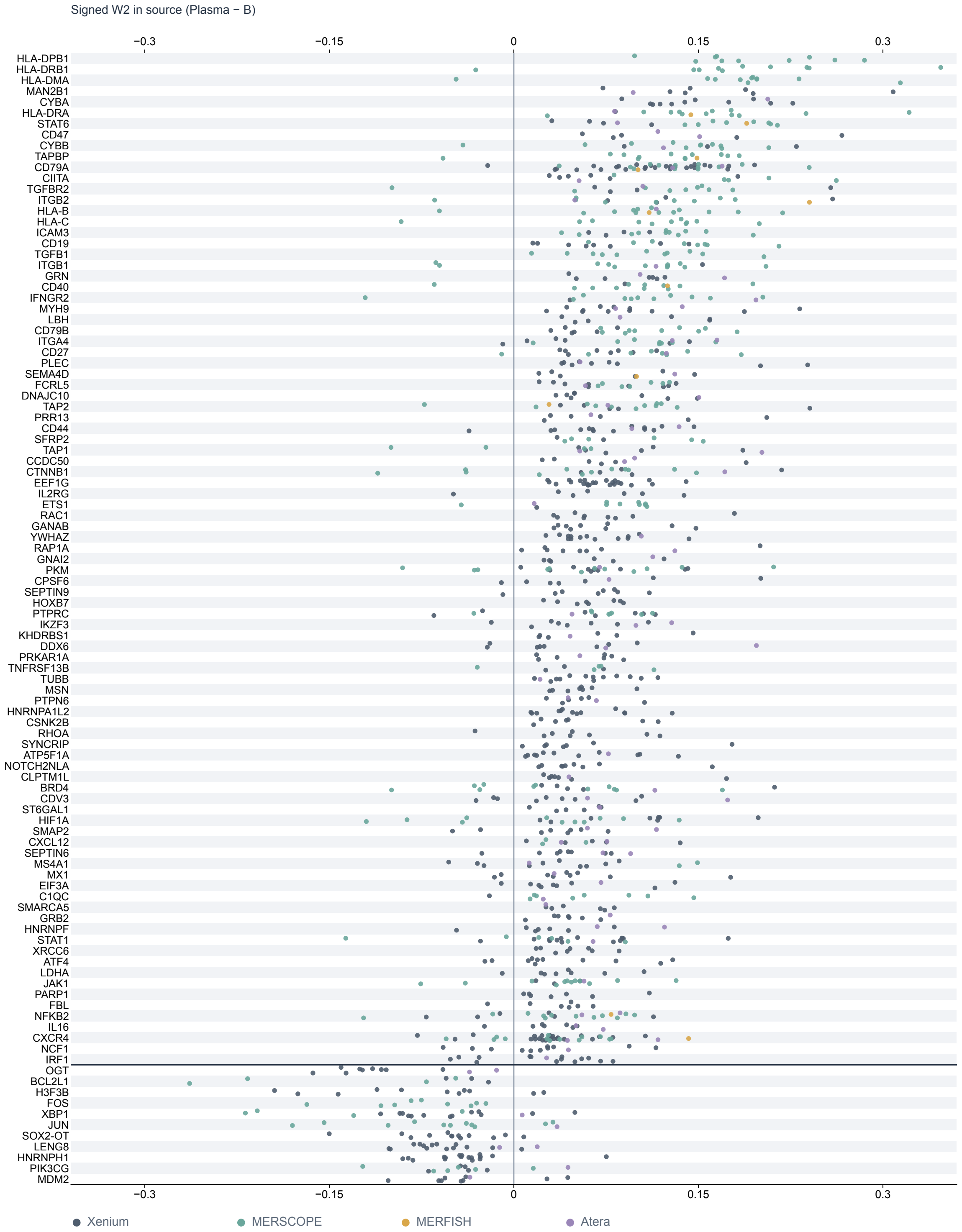


**Supplementary Figure 11  Radial shifts in each source.** Signed W2 (plasma − B) of each of the 104 detected genes (rows, outward genes above the line and inward genes below) in each source that tests it (dots). Colour, platform.

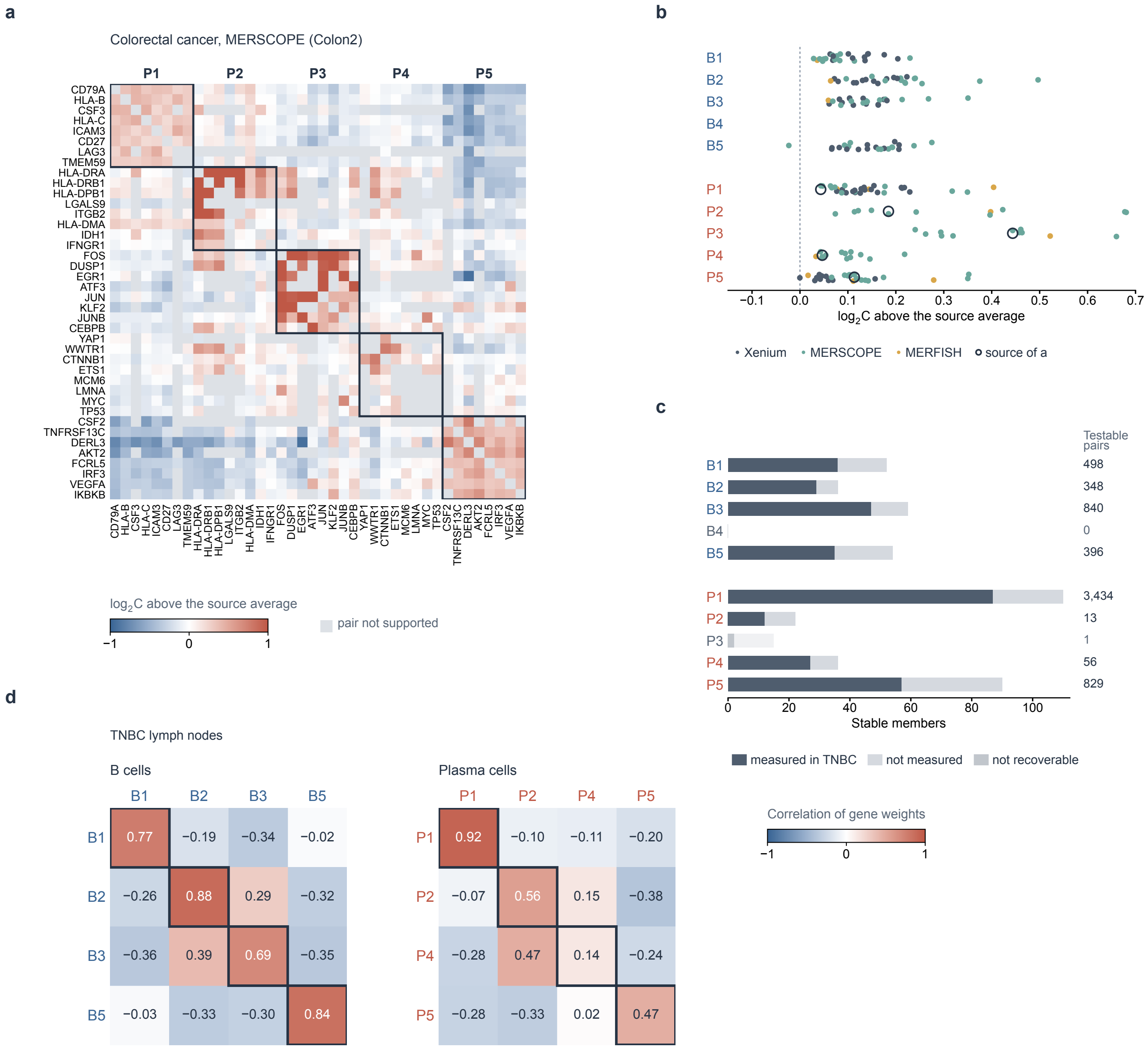


**Supplementary Figure 12 Co-occurrence programs within sources and in held-out TNBC lymph nodes.** **a**, Co-occurrence in plasma cells of the colorectal cancer MERSCOPE source Colon2 for the eight top-ranked genes of each of the five programs on its panel, each gene in the program of its largest share as in Fig. 3d. Colour, $\log_2 C$ above the mean of all supported pairs in the source. Boxes, programs. Grey, pair not supported. **b**, Co-occurrence of member pairs above that of all supported pairs for each program of B cells (B1–B5) and plasma cells (P1–P5) in each discovery source (dots). Colour, platform. Ring, the source in **a**. **c**, Stable members of each program measured (dark) or not measured (light) in the six held-out TNBC lymph-node sources, with the number of testable pairs among them (right). Faded, programs that cannot be recovered on the TNBC panel. **d**, Correlation over genes between the weights of TNBC genes placed on the fixed programs of Fig. 3d (columns) and their discovery weights (rows), for the recoverable programs of B cells (left) and plasma cells (right). Numbers, correlation. Bold boxes, $q < 0.05$.

**Supplementary Table 1 External RNA-level endoplasmic reticulum (ER) measurements for genes of program P1.** MERFISH with KDEL ER immunostaining in U-2 OS cells (ER versus non-ER cytoplasm; ER-enriched at $\log_2$ fold change $> 0$ and Bonferroni-adjusted $P < 10^{-10}$, the thresholds of the original study; Xia et al., PNAS 2019) and RNA-seq of ER-bound ribosomes labelled by LRRC59 or SEC61$\beta$ BioID in HEK293T cells (versus mock purification; lists filtered by the original study at $\geq$ 2-fold and adjusted $P <$ 0.05; Hoffman et al., J. Biol. Chem. 2019). The other 11 of the 20 highest-weight P1 genes were not measured in the first study and not listed in the second.

| Gene | Study | Cells | Measurement | $\log_2$ FC | Adjusted $P$ | Classification |
|---|---|---|---|---|---|---|
| *RPN1* | Xia et al. 2019 | U-2 OS | ER vs non-ER | 1.95 | $1.0 \times 10^{-220}$ | ER-enriched |
| *PDIA4* | Xia et al. 2019 | U-2 OS | ER vs non-ER | 2.41 | $1.0 \times 10^{-218}$ | ER-enriched |
| *CKAP4* | Xia et al. 2019 | U-2 OS | ER vs non-ER | 1.78 | $4.4 \times 10^{-218}$ | ER-enriched |
| *NUCB2* | Xia et al. 2019 | U-2 OS | ER vs non-ER | 1.05 | $1.2 \times 10^{-32}$ | ER-enriched |
| *SLAMF7* | Xia et al. 2019 | U-2 OS | ER vs non-ER | 0.81 | $3.5 \times 10^{-4}$ | below threshold |
| *TMEM59* | Xia et al. 2019 | U-2 OS | ER vs non-ER | 0.49 | 1 | below threshold |
| *ICAM3* | Xia et al. 2019 | U-2 OS | ER vs non-ER | $-0.09$ | 1 | below threshold |
| *HLA-B* | Hoffman et al. 2019 | HEK293T | LRRC59 ribosomes | 1.23 | $3.4 \times 10^{-6}$ | enriched |
| *HLA-B* | Hoffman et al. 2019 | HEK293T | SEC61$\beta$ ribosomes | 1.05 | $1.7 \times 10^{-4}$ | enriched |
| *HYOU1* | Hoffman et al. 2019 | HEK293T | LRRC59 ribosomes | 1.21 | $3.8 \times 10^{-7}$ | enriched |

**Supplementary Table 2 Sensitivity analyses of the B and plasma cell results** (separate Excel file). Each sheet gives one analysis. R_reruns and R_reruns_counts: the radial test rerun without the sources of each cancer type, of each platform or of the TNBC lymph nodes, and with only one group of sources. C_reruns and C_reruns_counts: the co-occurrence test rerun without each cancer type or with one group of sources only. Isolated_R and Isolated_C: MHC-II genes and pairs in each set of cells without MHC-II-high neighbours. Program_refits: programs refitted without one cancer type or with one group of sources only. Program_support: support of every program in each source. TNBC_supervised: the genes of the held-out TNBC lymph nodes placed on the fixed programs (Supplementary Fig. 12d). TNBC_projection and TNBC_refit: supportive readouts, the projection of each program onto the TNBC lymph nodes and the programs fitted independently there. Expression_GSEA: Hallmark enrichment in isolated cells, from the cell interior and in each source. The README sheet describes each sheet.
